# Genome annotation of *Caenorhabditis briggsae* by TEC-RED identifies new exons, paralogs, and conserved and novel operons

**DOI:** 10.1101/2021.09.24.461604

**Authors:** Nikita Jhaveri, Wouter van den Berg, Byung Joon Hwang, Hans-Michael Muller, Paul W. Sternberg, Bhagwati P. Gupta

## Abstract

The nematode *Caenorhabditis briggsae* is routinely used in comparative and evolutionary studies involving its well-known cousin *C. elegans*. The *C. briggsae* genome sequence has accelerated research by facilitating the generation of new resources, tools, and functional studies of genes. While substantial progress has been made in predicting genes and start sites, experimental evidence is still lacking in many cases. Here, we report an improved annotation of the *C. briggsae* genome using the *Trans*-spliced Exon Coupled RNA End Determination (TEC-RED) technique. In addition to identifying the 5’ ends of expressed genes, we have discovered operons and paralogs. In summary, our analysis yielded 10,243 unique 5’ end sequence tags with matches in the *C. briggsae* genome. Of these, 6,395 were found to represent 4,252 unique genes along with 362 paralogs and 52 previously unknown exons. These genes included 14 that are exclusively trans-spliced in *C. briggsae* when compared with *C. elegans* orthologs. A major contribution of this study is the identification of 493 operons, of which two-thirds are fully supported by tags. In addition, two SL1-type operons were discovered. Interestingly, comparisons with *C. elegans* showed that only 40% of operons are conserved. Of the remaining operons, 73 are novel, including 12 that entirely lack orthologs in *C. elegans*. Further analysis revealed that four of the 12 novel operons are conserved in *C. nigoni.* Altogether, the work described here has significantly advanced our understanding of the *C. briggsae* system and serves as a rich resource to aid biological studies involving this species.

## INTRODUCTION

Nematodes are a mainstay in fundamental biological research. While most work has been based on *C. elegans* over the last half a century since its proposed role as a model organism (Brenner 1974), the close relative *C. briggsae* offers many of the same advantages in carrying out studies. Despite diverging roughly 20-30 million years ago (Cutter 2008), the two species exhibit similar behavioral, developmental, and morphological processes including a hermaphroditic mode of reproduction (Gupta *et al*. 2007). Moreover, many of the experimental techniques and protocols developed to manipulate *C. elegans* can be adopted to *C. briggsae* with minimal to no modification (Baird and Chamberlin 2006; Gupta *et al*. 2007). These features make *C. briggsae* - *C. elegans* an ideal pair for comparative and evolutionary studies.

The genome of *C. briggsae* was sequenced many years ago and revealed extensive genomic and genic conservation with *C. elegans* (Stein *et al*. 2003). Subsequent work reported the assembly of genomic fragments into chromosomes and improved gene predictions (Hillier *et al*. 2007; Ross *et al*. 2011). While a diverse array of techniques have been applied to improve the annotation of the *C. elegans* genome (Hwang *et al*. 2004; Spieth and Lawson 2006; Hillier *et al*. 2009; Salehi-Ashtiani *et al*. 2009; Allen *et al*. 2011), a similar approach is lacking for *C. briggsae*. The current *C. briggsae* genome annotation is largely based on homology with the *C. elegans* genome. More analysis that uses experimental data gathered directly from *C. briggsae* is needed to improve gene identification and gene models. To this end, we used *trans*-spliced exon coupled RNA end determination (TEC-RED) (Hwang *et al*. 2004), a technique based on exploiting the phenomenon of spliced leader (SL) *trans*-splicing which has been observed in nematodes and several other phyla including platyhelminths, chordates and trypanosomes (Lasda and Blumenthal 2011).

The advantage of TEC-RED compared to other genome annotation techniques like EST (Marra *et al*. 1998) and SAGE (Velculescu *et al*. 1995) is that it is capable of identifying transcripts of most expressed genes, and uniquely allows for the identification of 5’ transcript start sites and alternative transcripts with different 5’ ends of a gene. The approach is based on two principles: one, a short sequence from the 5’ end of a transcript can be used to uniquely identify the initiation site of the transcript, and two, the 5’ ends of mRNAs are spliced to common leader sequences known as spliced leader (SL) sequences. The SL *trans*-splicing process involves replacing the outron of a pre-mRNA with a 22 nucleotide SL sequence donated by a 100-nucleotide small ribonucleoprotein (snRNP) (Blumenthal 2005; Allen *et al*. 2011). *C. elegans* and *C. briggsae* both have two types of spliced leader sequences: SL1 and SL2 (Qian and Zhang 2008; Blumenthal *et al*. 2015).

We recovered well over 120,000 5’ end tags from sequencing reactions representing 10,243 unique tags (7,234 for SL1; 3,009 for SL2) with matches in the *C. briggsae* genome. The tags were analyzed using WormBase release WS276 and it was found that more than 60% could be aligned to exons curated in WormBase (www.wormbase.org). Most of the tags were found to have unique hits in the genome and identified a total of 4,252 genes. Other tags with high confidence hits to unannotated regions or to multiple locations of the genome identified 52 novel exons and 362 paralog genes, respectively. The novel exons could either represent previously unknown genes or new exons of existing genes. The paralogs define 133 sets of two or more genes. Of these sets, 21 were confirmed as exact matches with known paralogs in WormBase. The rest could potentially be new paralogous pairs that need further validation. While the majority of the genes discovered by tags confirmed 5’ ends of genes listed in WormBase, there are many for which 5’ ends indicated by tags differ from current gene models, suggesting the need to revise existing annotations.

A comparison of the splicing pattern of *C. briggsae* genes with *C. elegans* revealed some changes. Specifically, 14 genes are spliced to leader sequences in *C. briggsae* but their *C. elegans* orthologs lack such splicing information. We also investigated the presence of operons. It was reported earlier that 96% of *C. elegans* operons are conserved in *C. briggsae* based on collinearity (Stein *et al*. 2003). Our analysis revealed a total of 1,199 operons including 493 for which splicing identities of two or more genes are reported in this study. Of these operons, 334 are fully supported by tags. Comparison of the latter with *C. elegans* revealed that 40% are conserved, the largest of which is composed of seven genes. Another 38% are termed partially conserved since gene sets do not fully correspond to any of the operons in *C. elegans*. The remaining are novel, i.e., consisting of divergent genes as well as genes whose *C. elegans* orthologs are not reported in operons. Of the divergent operons, four were found to be conserved in a closely related sister species, *C. nigoni*. Lastly, two SL1-type operons have been identified. Overall, the results presented in this study have substantially improved the annotation of the *C. briggsae* genome by identifying the 5’ ends of a large number of genes as well as discovering novel operons, new exons, and paralogs. The findings strengthen the utility of *C. briggsae* as a model organism and serve as a platform to accelerate comparative and evolutionary studies involving nematodes and other metazoans.

## MATERIALS AND METHODS

### Generation of tags

We followed the protocol described earlier for *C. elegans* (Hwang *et al*. 2004). Briefly, the steps involved purification of poly(A) RNA from the wild type *AF16* mixed stage animals, RT-PCR to generate cDNA, amplification of cDNAs using biotin-attached primers homologous to SL1 and SL2 sequences carrying mismatches to create *Bpm*I restriction enzyme site (Supplementary Tables 1-3), digestion of amplified cDNAs using *Bpm*I to produce short fragments (termed “5’ tags”), ligation of tags to adaptor DNA sequences, and sequential ligation of DNA to create concatenated products. The ligated DNA pieces were cloned into a vector and sequenced.

### 5′Tag sequence analysis and exon identification

We wrote several Perl scripts to analyze the tags and genes. A flowchart is provided in Supplementary figure 1. Briefly, tags were collected and assigned unique tag IDs. Tag locations in the genome were determined by comparing the tag sequence to WS176 and WS276 genome files, where orientation and chromosome location for each tag was noted. Subsequently the splice sequence for each tag was obtained by finding the seven bases directly upstream of each location where a tag matched on the genome.

The criteria to identify tag matches to exonic regions were described earlier (Hwang *et al*. 2004). These included ‘same orientation of the tag as that of the corresponding exon’, ‘distance to the first ATG’, ‘a minimum distance to the nearest in-frame stop codon’ and ‘presence of a splice acceptor sequence following the tag’. The latter was scored on how well they fit the consensus splice site ‘TTTTCAG’ (Blumenthal and Steward 1997). In cases where tags had multiple matches, we applied stricter splice acceptor site criteria. Perfect consensus sequence was given the highest weight. Sites having mismatches were assigned lower weights with priority given to conserved bases. While this approach resulted in most tags identifying unique exons, a small number still showed multiple matches and were used to search for potential paralogs (see below).

Each tag was used to find the nearest ATG of an open reading frame (ORF), i.e., the proposed start of a coding sequence (CDS). This ATG location was compared to known coordinates of start sites of nearest exons as annotated in WS176 and WS276 genome annotation (gff3) files. This was done using coordinates of annotated CDS. Two broad categories of exon matches were identified based on tags that had unique matches: one, where the 5’ end corresponded to the start of a known exon (first exon: 1a, internal exon: 1b) and two, matches for which the 5’ end differed from a nearest exon (Figure 1). Depending on the distance between the 5’ end and the exon, the second category of matches were further divided into two sub-categories. These consisted of exons that were either within 20 bp from the 5’ end (‘minor misprediction’) or further away (‘major misprediction’). The major misprediction sub-category also includes matches where 5’ ends were more than 3 kb away and may define brand new exons of existing genes as well as potentially new, previously unknown genes.

**Figure 1:**
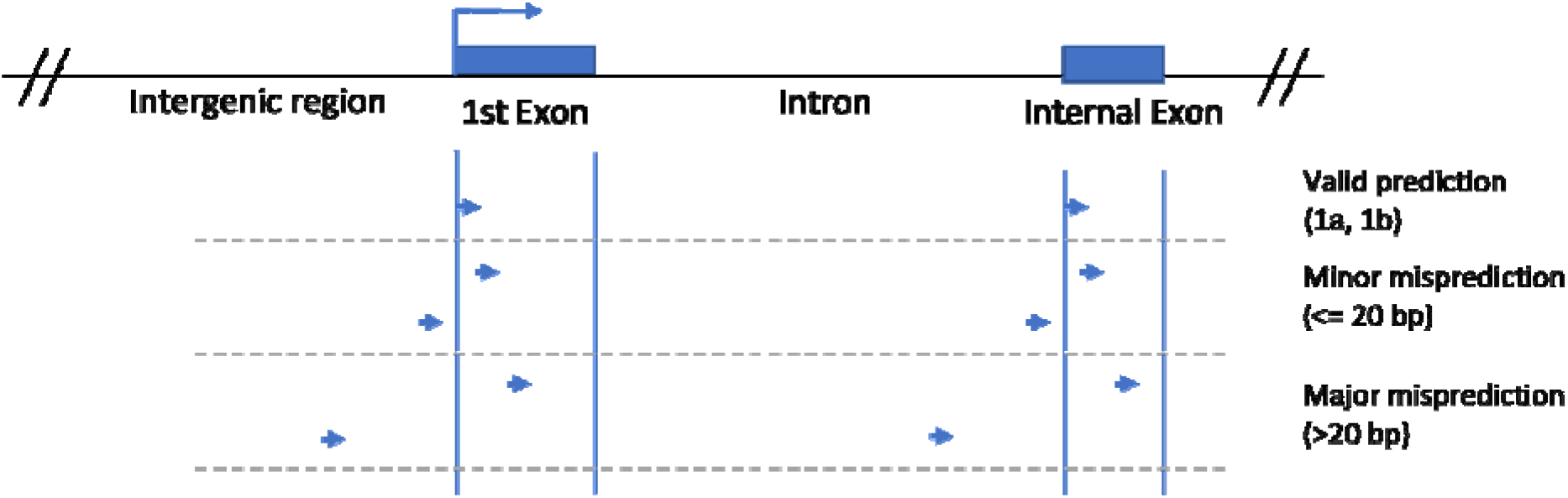
Representative model of locations of tag sequences within the genome. Three broad categories of matches are: valid prediction (termed 1a and 1b), minor misprediction, and major misprediction.

### Manual curation of genes

We found that 75 tag-matched genomic regions in the WS276 gff3 file had no known genes/exons within 3kb downstream of the matched ATG. The surrounding chromosomal regions of these matches were confirmed by manually searching the WormBase genome browser for presence of annotated exons. Of the 75, 21 were found to be false positives due to incorrect script calls. Two were excluded from analysis because the genes are not assigned to any chromosomes. The remaining 52 matches may represent novel exons.

### Analysis of intergenic regions and operons

The distance between two genes, termed ‘intergenic region’ (IGR) was determined based on the distance from the end of the 3’ UTR of an upstream gene to 5’ start of the coding sequence of the immediate downstream gene. Graphs were generated using Graphpad Prism 7.0 and Microsoft Excel. Genes having IGR >5000 bp (257) were excluded from the analysis. For pairs of genes where the second gene is located within the first gene, IGR length is calculated as a negative value. Intercistronic regions (ICRs) were calculated in the same manner. The ICR analysis was done only for genes identified to be part of an operon.

To identify genes that could be present in operons, all genes *trans*-spliced with SL2 or SL1/SL2 and present downstream of an SL1-spliced gene were categorized into a single operon model along with the upstream SL1 spliced gene. We based our assumption of genes being in an operon together on this pattern of splice leader sequences. If the splicing of the first upstream gene was unknown, the operon models were termed ‘non-tag-supported’ whereas those models in which the identity of the first upstream gene was known were termed ‘tag-supported’. We compared the ‘tag-supported’ operon models to those in *C. elegans* (WormBase) to determine how well operons are conserved. Based on the conservation of genes, the ‘tag-supported’ operons were classified into Exact match, Partial match, and Novel.

We examined the enrichment of germline genes in *C. briggsae* high confidence operons. For this, *C. elegans* orthologs were identified and researched for association with germline function (Wang *et al*. 2009). The significance of overlap was tested by the hypergeometric probability test. Next, to identify processes related to genes in operons, Gene Ontology (GO) (Ashburner *et al*. 2000) analysis was carried out for all operon genes. A similar analysis was conducted for genes present in *C. elegans* operons using a published data set (Allen *et al*. 2011).

### Paralog analysis

A total of 203 tags had multiple hits in the genome. Since many of these consisted of overlapping sequences, we retained only the longest tags. This filtering step narrowed down the count to 158. The genes identified by these tags were compared with annotated paralogs in WormBase. The matches allowed us to place the predicted paralogs into three different categories.

### Uniquely spliced C. briggsae genes

To identify the genes that are trans splicing in *C. briggsae* but not in *C. elegans*, we used datasets published by two different groups that together constitute the most complete collection of *trans*-spliced genes in *C. elegans* (Allen *et al*. 2011; Tourasse *et al*. 2017). Initial comparisons with the Allen et al. dataset revealed 198 genes that are present only in our analysis. The number was further reduced to 14 genes when compared with the Tourasse et al. study (Supplementary data file 3).

## RESULTS

### Overview of the TEC-RED method in *C. briggsae*

To implement the TEC-RED approach to identify transcripts, we first isolated *C. briggsae* mRNAs containing an SL1 or SL2 sequence at their 5′ ends. A total of 121,189 5′ tags (91,733 for mRNA with an SL1 and 29,456 for mRNA with an SL2 spliced leader sequence) were recovered from DNA sequencing reactions. These tags represent 14,678 different sequences, of which 10,400 (71%) are for SL1 and 4,278 (29%) for SL2 sites. More than two-thirds of all tags found matches in the genome (10,243, 70%), of which 46% are unique, i.e., matching only once and others matching multiple times (Table 1). The proportions were similar for both spliced leader categories, demonstrating no bias in the experimental protocol. The remaining 4,435 tags (30%) had no match, likely due to reasons such as sequencing errors, gaps in the genome sequence, and incorrect sequence assembly.

**Table 1:** Overview of SL1 and SL2 5’ tags identified in the study.

|  | **Total unique tags** | **Matches in genome** | **Unique hits** | **Multiple hits** |
| --- | --- | --- | --- | --- |
| All | 14,678 | 10,243 | 4,753 | 5,490 |
| SL1 | 10,400 | 7,234 | 3,281 | 3,953 |
| SL2 | □ 4,278 | 3,009 | 1,472 | 1,537 |

### Exon validations and predictions in *C. briggsae* based on 5’ tag matches

After filtering the matches (see Methods), 62.5% of all tags (6,395 of 10,243) were retained for further analysis (Table 2). Next, we determined the locations of these tags relative to annotated exons in WormBase. Most of the tags (6,192, 96.8%) matched uniquely to one exon, with a small number having multiple matches (203, 3.2%) (Supplementary data file 1). For both SL1 and SL2 tags, roughly 80% of the matches correspond to known 5’ ends of annotated genes (Category 1a), providing support to existing gene models in WormBase. Less than one percent of the tags matched to internal exons (Category 1b), suggesting an alternate 5’ end of the corresponding genes. The remaining tags identified start sites that differed from current WormBase gene models and were categorized as mispredicted genes. In most of these cases (roughly three-quarters of all mispredictions) the nearest exon was more than 20 bp away. This leads us to suggest that, particularly in these cases, existing gene models may need to be revised. These exons may define new 5’ ends of known genes as well as novel, previously unidentified genes. More experiments are needed to investigate these possibilities. As expected, both types of tags, i.e., with unique and multiple hits have a similar distribution of categories (Figure 2, Supplementary data file 1).

**Table 2:**
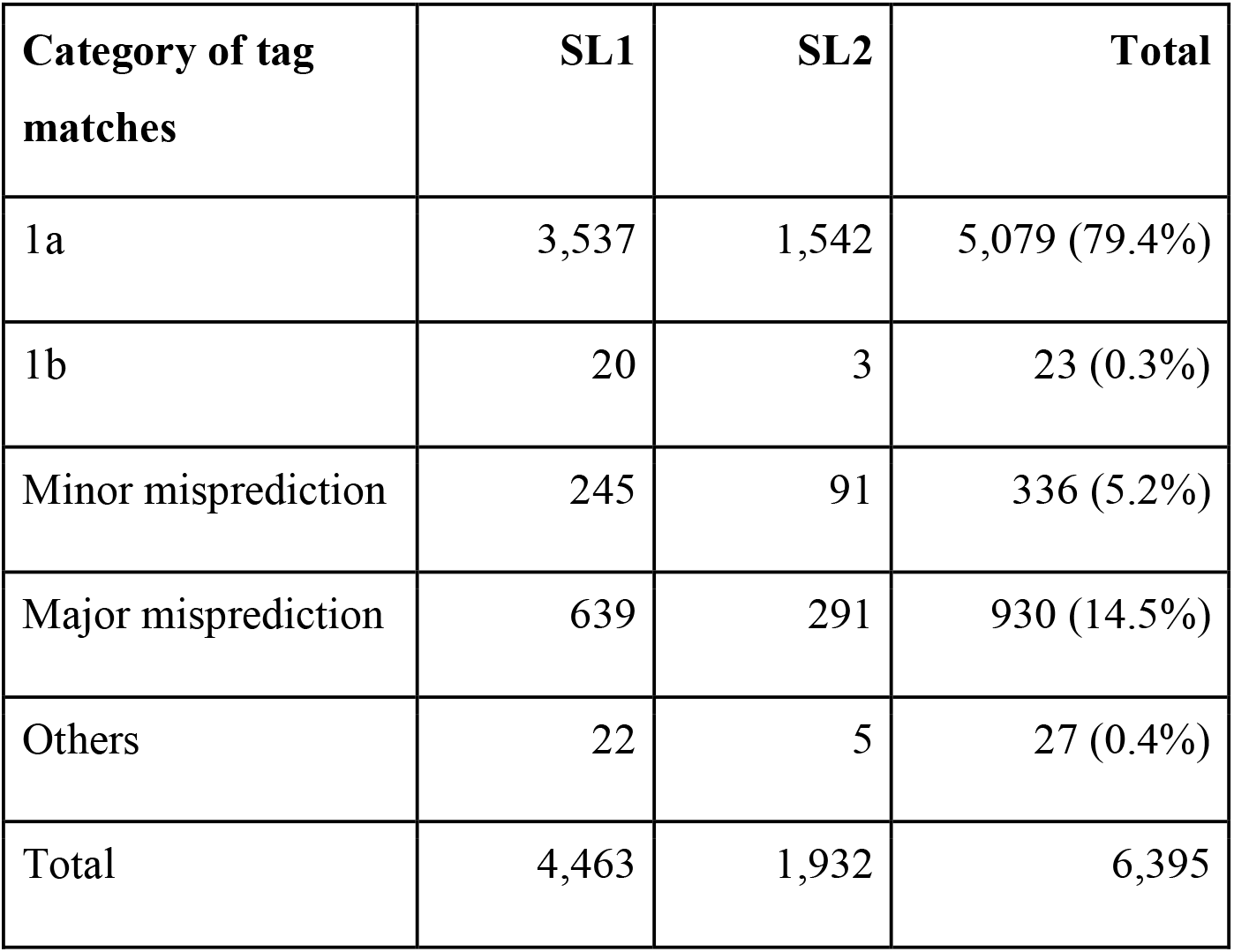
Breakdown of tag matches into different categories. The numbers include both unique and multiple hits. Tag matches termed as ‘Others’ are those that cannot be placed uniquely into any of the main categories.

| Category of tag matches | SL1 | SL2 | Total |
| --- | --- | --- | --- |
| 1a | 3,537 | 1,542 | 5,079 (79.4%) |
| 1b | 20 | 3 | 23 (0.3%) |
| Minor misprediction | 245 | 91 | 336 (5.2%) |
| Major misprediction | 639 | 291 | 930 (14.5%) |
| Others | 22 | 5 | 27 (0.4%) |
| Total | 4,463 | 1,932 | 6,395 |

**Figure 2.**
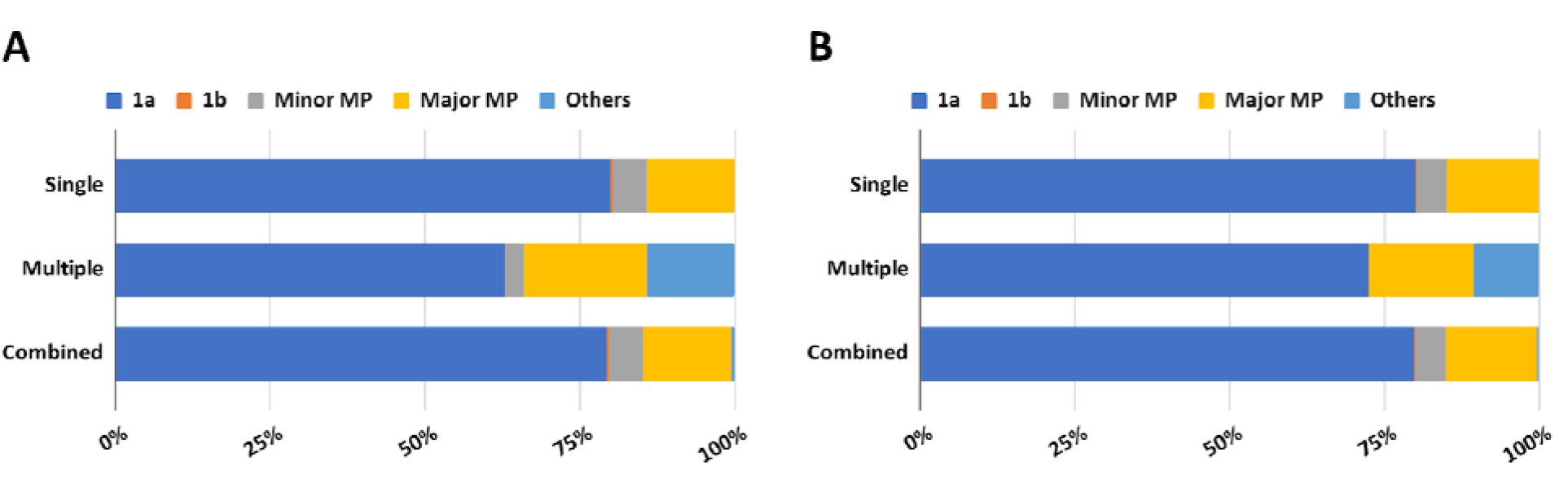
Proportion of tags belonging to different categories. The majority of SL1 **(A)** and SL2 tags **(B)** have single (unique) hits in the genome and belong to category 1a, i.e., predicted 5’ ends match with WormBase gene models. Minor MP: Minor misprediction, Major MP: Major misprediction, Others: mix category of matches.

### Identification of genes based on tag matches

Next, we compiled a list of *C. briggsae* genes based on exons identified by unique tags. A total of 4,252 genes were recovered by SL1 and SL2 tags (Supplementary data file 2). Almost two-thirds of the genes (65%) are spliced with SL1 and 18% with SL2. Another 18% of exons matched with both SL1 and SL2 tags (SL1/SL2), suggesting the genes are part of hybrid operons (Allen et al., 2011) (Table 3, Supplementary data file 2). Based on their genomic locations, these genes are roughly evenly distributed on the chromosomes except for Chr. X which had the lowest gene count. However, the trend was different for gene density with Chr. III being the densest chromosome and Chr. X the sparsest (Supplementary table 4). Whether the uneven distribution is by chance or a characteristic of *trans*-spliced genes in *C. briggsae* remains to be seen. A tiny fraction of genes (0.1%) is located in unmapped genomic fragments.

**Table 3:** Breakdown of genes by spliced leader sequences.

|  | Number of genes |
| --- | --- |
| SL1 type | 2,750 (65%) |
| SL2 type | 743 (18%) |
| SL1/SL2 type | 759 (18%) |
| **Total** | 4,252 |

**Table 4.** Genes supported by the presence of a single 5’ end (single transcript). Numbers refer to genes identified by SL1, SL2 and SL1/SL2 tags. The genes have been divided further into various categories based on distance from the nearest exon (see figure 1). Novel exons and potential paralogs are excluded.

|  | **All** | **SL1** | **SL2** | **SL1/SL2** |
| --- | --- | --- | --- | --- |
| Matching first exon (1a) | 3,288 | 2,143 (65.2%) | 558 (17.0%) | 587 (17.9%) |
| Matching internal exon (1b) | 16 | 14 (87.5%) | 1 (6.2%) | 1 (6.2%) |
| Minor misprediction of first or internal exon | 204 | 146 (71.6%) | 34 (16.7%) | 24 (11.7%) |
| Major misprediction of first or internal exon | 517 | 357 (69%) | 127 (24.6%) | 33 (6.4%) |
| **Total** | **4,025** | **2,660** | **720** | **645** |

The recovery of *C. briggsae* genes prompted us to examine evolutionary changes in *trans*-splicing. A comparison with *C. elegans* studies (Allen *et al*. 2011; Tourasse *et al*. 2017) revealed 14 genes that appear to be uniquely spliced to leader sequences in *C. briggsae* but not in *C. elegans* (Supplementary data file 3).

Next, we searched for transcripts resulting from *cis*-splicing of the *C. briggsae* genes. Almost 95% of the curated genes (4,025 of 4,252) were found to be associated with unique tag sequences, i.e., 5’ ends matched to just one exon, providing support for the presence of a single transcript for these genes (Table 4). In the majority of cases (82%, 3,288 of 4,025), the tag-identified 5’ ends matched with a known first exon (category 1a tags). Less than one percent of the tags identify 5’ ends that match with internal exons (category 1b). The remaining genes (18%) consist of exons belonging to minor and major misprediction categories. The rest of the genes (5%, 227 of 4,252) identified by tags consist of those that produce multiple transcripts (Table 5). In 84% of these cases, at least one 5’ end identified by tags matched with the first exon (category 1a). Five of the genes were alternatively spliced using internal exons as the 5’ start site (category 1b). Most of the genes consisted of at least one major mispredicted exon, suggesting that genes with multiple splice variants require further validation.

**Table 5:** Genes supported by the presence of multiple 5’ ends. Numbers refer to genes identified by SL1, SL2 and SL1 and SL2 tags. These genes have been divided further into various categories based on distance from the nearest exon (see figure 1). Genes for which exons belong to multiple categories are grouped as ‘Others’. Novel exons and potential paralogs are excluded.

|  | **All** | **SL1** | **SL2** | **SL1/SL2** |
| --- | --- | --- | --- | --- |
| Matching first exon (1a) and matching internal exon (1b) | 5 | 2 (25%) | 0 | 3 (75%) |
| Matching internal exons (1b) | 0 | 0 | 0 | 0 |
| Matching first exon (1a) and minor misprediction of one or more internal exons | 39 | 14 (36%) | 7 (18%) | 18 (46%) |
| Matching first exon (1a) and major misprediction of one or more internal exons | 145 | 57 (39%) | 14 (10%) | 74 (51%) |
| Others | 6 | 2 (33%) | 0 | 4 (67%) |
| All mispredicted exons (minor and major) | 32 | 16 (50%) | 2 (6%) | 14 (44%) |
| **Total** | **227** | **91** | **23** | **113** |

As mentioned above, 203 tags had multiple matches in the genome. Further analysis narrowed down the set to 158 unique sequences (see Methods). We reasoned that these tags may represent paralogs and performed searches in WormBase. The analysis identified 133 potential paralog sets consisting of 362 genes. These sets fall into three distinct categories (Supplementary data file 10). Paralogs that fully matched with WormBase annotation were termed ‘Exact Match’ (21 paralogous sets, 42 genes). The other sets matched only partially or did not match to paralog sets recorded on WormBase (Partial Match: 66 sets, 174 genes; No Match: 46 sets, 146 genes). It is worth mentioning that about half of the genes in the No Match category have no paralogous information available, whereas the remaining half have paralogs in WormBase but these did not match with our analysis. To further validate the paralogous relationships, we determined chromosomal locations of the genes. Gene duplications arising from mechanisms such as slipped-strand mispairing can cause the creation of paralogous genes in adjacent stretches of sequence on the same chromosome (Levinson and Gutman 1987). In *C. elegans*, paralogs originating from gene duplications are more than twice as likely to be present on the same chromosome and tend to be located closely together (Semple and Wolfe 1999). Additionally, studies in humans and other higher eukaryotes have revealed that intergenic distances between paralogous genes are smaller than random gene pairs on the same chromosome (Ibn-Salem *et al*. 2017). Of the paralog sets identified in this study, 63% (84 sets) were present on the same chromosome including 35% (16 sets) that belong to the No Match category. The IGR analysis revealed that the distances in five cases are less than 10 kb (Supplementary table 5), which is more than 500-fold shorter than the average distance between a random pair of genes on the same chromosome (5.58 +/- 0.89 Mb in *C. elegans*) (Lee and Sonnhammer 2003).

### Validations of TEC-RED-identified transcripts

We took three different approaches to validate subsets of TEC-RED predictions with the goal of demonstrating the usefulness of the technique in improving gene identification and gene models. One approach involved comparing different categories of tag-identified exons between two WormBase releases. As described above, a significant number of exons are categorized as minor and major mispredictions (22%, 943 of 4,252; see Tables 4 and 5). We hypothesized that mispredicted exons may be confirmed with improvements in genome annotation. To test this hypothesis, 1a category of transcripts were compared with those reported in an older WormBase release (WS176). The analysis involved SL1 spliced transcripts belonging to category 1a (2,143) (Table 4). As expected, a vast majority of the genes (74%, 1,583) are in category 1a in both releases, providing support for these gene models (Figure 3A, 3D, Supplementary data file 4). The next two largest categories consist of genes that are mispredicted (11.7%, 250 genes) and newly predicted, i.e., absent in WS176 (13.2%, 286 genes). Few genes (0.5%, 11) have start sites that correctly match with internal 5’ ends of internal exons. The rest (0.5%, 14 genes) could not be uniquely placed into a single category since these had multiple tag matches in the older annotation. Roughly similar results were obtained by analyzing 1a category of SL2 spliced and SL1/SL2 spliced genes (182 genes, 115 genes, respectively) (Table 4; Figure 3B-D; Supplementary data file 4). Altogether, 858 annotation improvements are supported by our analysis. The demonstrated improvements in gene identification and genome annotation as observed in WS276 prove the accuracy of our 5’ start site determination method.

**Figure 3:**
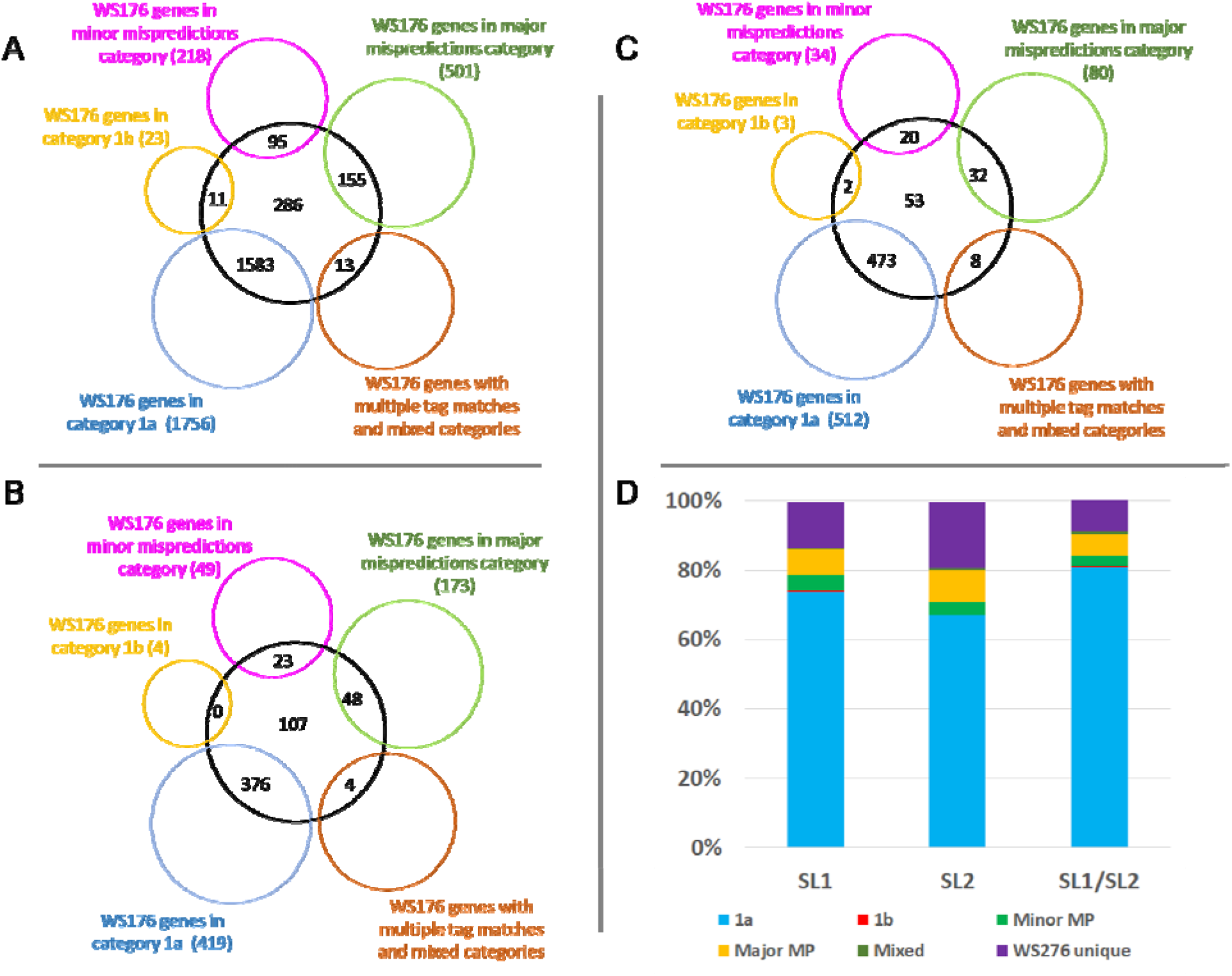
Reclassification of genes from WS176 categories to Category 1a in WS276. Only single transcript genes were compared. **(A-C)** Venn diagrams, with WS276 genes of category 1a in black circles and WS176 genes of various categories in colored circles. Numbers in overlapping circles represent genes of a given category in WS176 that are annotated as 1a type in WS276. Numbers in the middle of black circles (non-overlapping) represent genes that are unique to WS276 analysis (**A**, 286 or 13.2% of SL1-spliced; **B**, 107 or 19.0% of SL2-spliced; **C**, 53 or 9% of SL1/SL2 hybrid-spliced) whereas those in brackets next to colored circles are total genes identified by tag searches in WS176. **(D)** Histogram showing the proportion of genes with matching 5’ ends in WS276 (category 1a) that overlap with various categories in the WS176 analysis.

The second type of validation focused on a subset of the major misprediction category of genes whose 5’ ends mapped more than 3 kb away from nearest exons. Most of these (94%, 49 of 52) are in intergenic regions (Supplementary data file 5). 37% (19 of 52) of the exons are supported by RNA sequencing reads (WormBase), providing proof of accuracy to our method (Supplementary figure 2). These novel exons are likely to either belong to nearby existing genes or define brand new genes.

The last set of validations consisted of comparisons with *C. elegans* gene models. In this case category 1b of single and multiple transcripts (Tables 4 and 5 respectively) were manually examined. The results showed that 38% of newly discovered 5’ ends (6 single transcript and two multiple transcripts) are supported by *C. elegans* orthologs (Supplementary figure 3, Supplementary data file 6), providing further support to our analysis. We took a similar approach to analyze a subset of transcripts in the major mispredictions category. Of the 10% of such predictions that were tested, 34% (17 of 50) are supported by WormBase *C. elegans* gene models. With this success rate, another 115 of the remaining single transcript genes of the major misprediction category are likely to be validated. Overall, the 5’ tag analysis serves as a rich resource to improve the *C. briggsae* genome annotation.

### Discovery of operons

The identification of genes based on unique tag matches in *C. briggsae* allowed us to search for operons. In *C. elegans* it has been shown that the first gene in an operon is SL1 spliced (Conrad *et al*. 1991), whereas downstream genes are spliced either with SL2, SL2 variants or both SL1 and SL2 (Blumenthal 2005). Genes that are both SL1 and SL2 spliced cause the operon they are part of to be considered as ‘hybrid operon’. Ultimately, global analysis of *trans*-splicing in *C. briggsae* will reveal all operons and operon genes.

Our data suggests the existence of up to 1,199 *C. briggsae* operons (Table 6, Supplementary data file 7). These include 334 operons that are fully supported by tags, i.e., we were able to determine the splicing pattern of every gene, with operons ranging from two to seven genes The remaining 865 operons (ranging between two to six genes) are categorized as ‘Predicted operons’ since the splicing identity of the first gene in these cases remains to be determined. In this set, the predicted operons that contain three or more genes (159) are large enough to be labeled as bona fide operons. Added together with the 334 fully supported operons, this allows us to report at least 493 operons in *C. briggsae* with sufficient certainty.

**Table 6:** Breakdown of *C. briggsae* operons based on the number of genes present. Operons are placed into two broad categories, those consisting entirely of genes with known spliced leader sequences (Tag-supported) and others where the splice leader identity of the first gene is unknown (Predicted).

|  | No. of operons | Operons consisting of |  |  |  |  |  |
| --- | --- | --- | --- | --- | --- | --- | --- |
|  |  | 2 genes | 3 genes | 4 genes | 5 genes | 6 genes | 7 genes |
| Tag-supported operons | 334 | 263 | 54 | 14 | 2 | 0 | 1 |
| Predicted operons | 865 | 706 | 125 | 26 | 7 | 1 | 0 |

In *C. elegans*, operon genes tend to be very closely spaced, typically having an ICR of less than 1 kb (Allen *et al*. 2011; Blumenthal *et al*. 2015). To examine whether the same is true in *C. briggsae*, we calculated ICRs and found that a vast majority of the genes (78%) are separated by less than 200 bp (Figure 4). We also determined intergenic distances (IGRs) for SL1 and SL1/SL2 hybrid spliced genes discovered in our study. The results suggested that the IGR to the nearest gene upstream of SL2-spliced genes is smaller compared to those spliced with SL1 and SL1/SL2. While the SL2-spliced genes have a median distance of 180 bp, the medians of SL1 and SL1/SL2 spliced genes are 4,631 bp and 1,242 bp, respectively (Figure 5A). Furthermore, as we would expect, genes with larger IGRs are more likely spliced with SL1 than SL2 or SL1/SL2 (Figure 5B, Supplementary data file 8) and are thus less likely to be part of the same operon.

**Figure 4:**
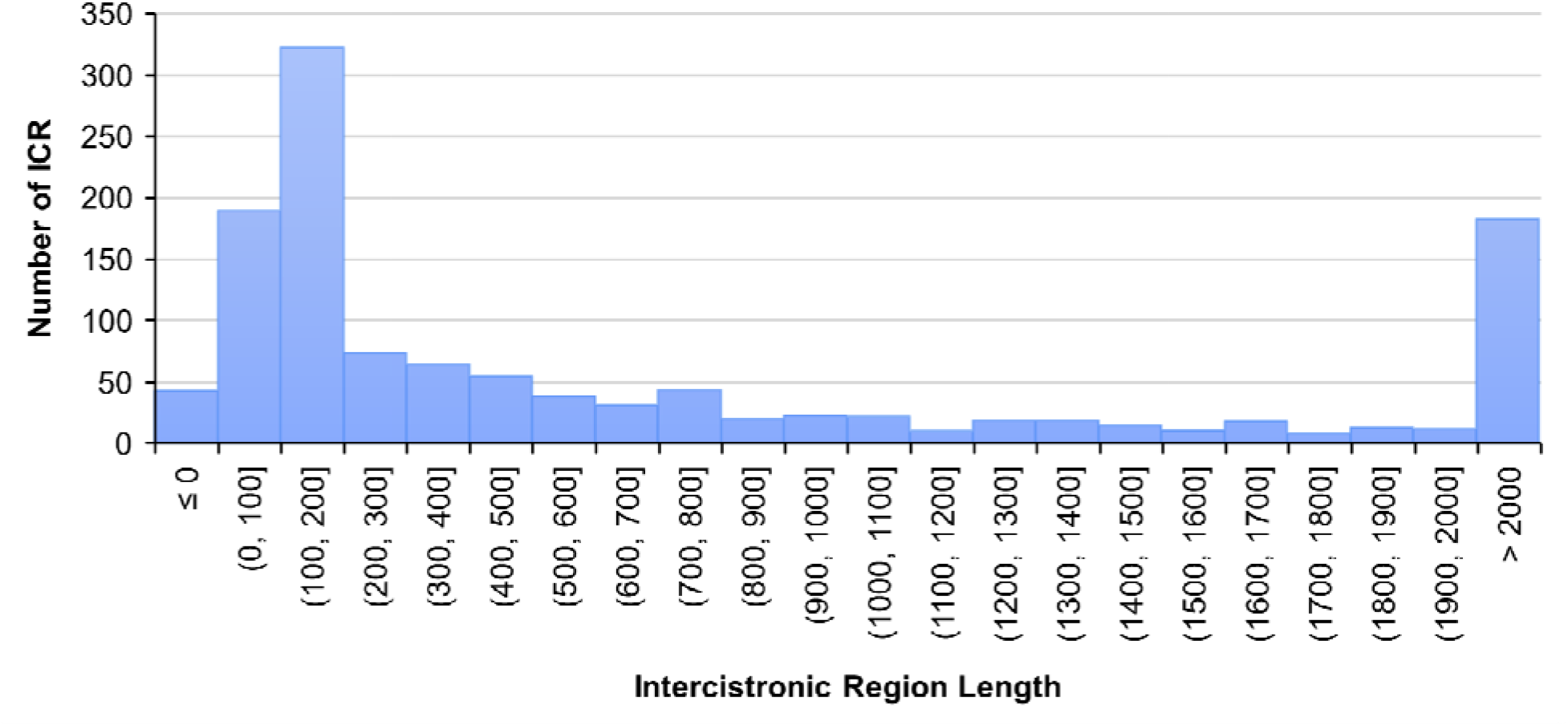
Frequencies of ICR lengths between SL2 and hybrid-spliced genes in operons. ICRs are sorted in bins of 100 nucleotides. For pairs of genes where the second gene is within the first gene, ICR is calculated as a negative value. For bin sizes, round brackets indicat exclusive bound, square brackets indicate inclusive bounds. Genes with larger than 2kb ICRs are shown as a single peak.

**Figure 5:**
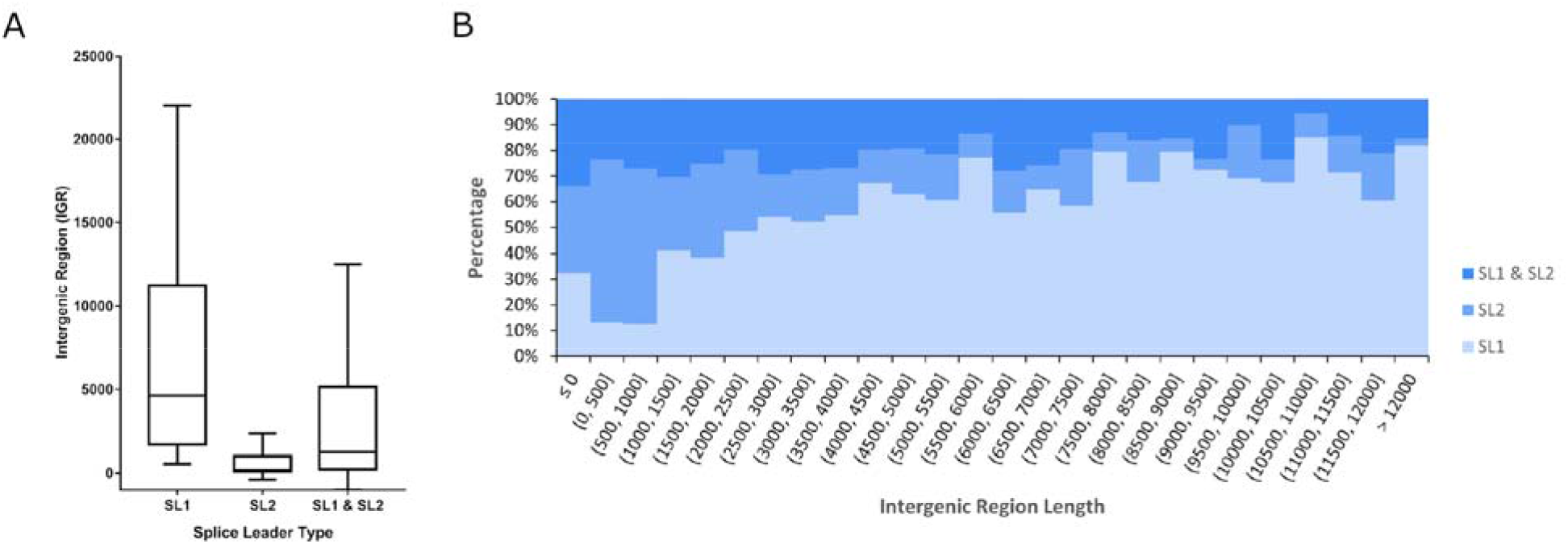
A: IGRs of genes identified by tag matches. (A) Box plots show IGRs for SL1-spliced, SL2-spliced, and SL1/SL2-spliced genes. The inside line marks the median, lower and upper lines represent the borders of the 25th and 75th quartile of the data sample, respectively. Whiskers enclose the 10-90% range of the data. **(B)** 100% stacked columns of IGR length. Lengths are sorted in bins of 500 nucleotides. For pairs of genes where the second gene is overlapping or inside the first gene, length was calculated as a negative value. For bin sizes, round brackets indicate exclusive bound, square brackets indicate inclusive bounds.

### Tag-supported operons

We examined the conservation of tag-supported operons in *C. elegans*. The analysis of orthologs helped define three distinct categories (Supplementary data file 7). The two largest categories are named ‘exact match’ and ‘partial match’ operons (40% and 38%, respectively). Exact match operons consist entirely of *C. elegans* orthologs, whereas in partial match operons only some of the genes are conserved. The remaining one-fifth of operons define a third category, named ‘novel’ (73). While a majority of these (61, 18%) consist of conserved genes whose orthologs are not present in *C. elegans* operons, others (12, 4%) consist of divergent, *C. briggsae*-specific genes.

The largest *C. briggsae* operons (CBROPX0001) consists of seven genes, six of which (*CBG25571, CBG03062, CBG25572, CBG03061, CBG03060, CBG03059*) are conserved in *C. elegans* and are part of the orthologous operon CEOP2496. The 5th gene in CBROPX0001 (*CBG25573*) does not appear to have a *C. elegans* ortholog. Syntenic alignments revealed that *CBG25573* is conserved in *C. brenneri*, suggesting that the gene may have been lost in the *C. elegans* lineage (Supplementary figure 4). While we did not recover a tag for *Cbr-rpb-6 (CBG03063)*, whose ortholog is the first gene in CEOP2496, we hypothesize that it is part of *C. briggsae* operon CBROPX0001 based on the distance from its neighbor *CBG25571* (195 bp) (Figure 6). More experiments are needed to confirm if *Cbr-rpb-6* is the eighth gene in CBROPX0001.

**Figure 6.**
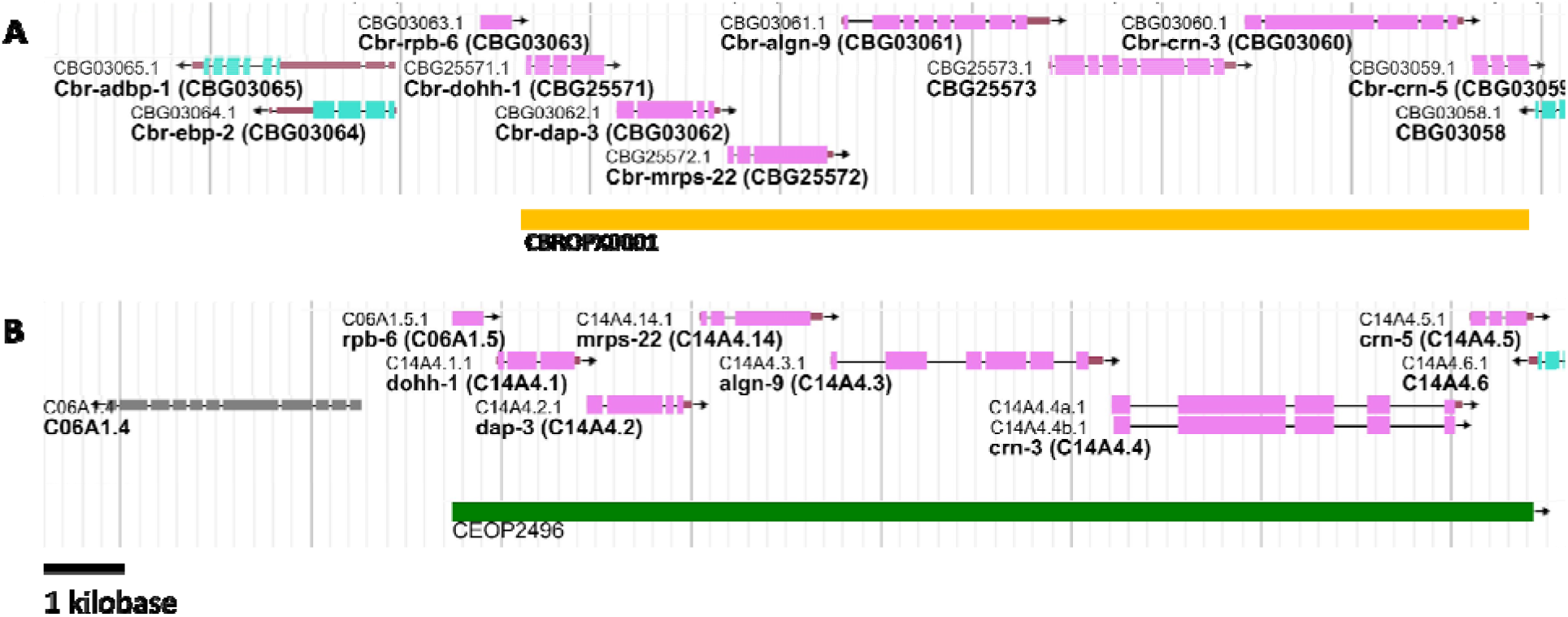
Genomic regions of *C. briggsae* CBROPX0001 and *C. elegans* CEOP2496. **(A)** CBROPX0001 is proposed to contain at least seven, and possibly eight, genes depending on the inclusion of *CBG03063*. **(B)** Homologous *C. elegans* operon CEOP2496 contains seven genes. Genomic feature visualizations in this and subsequent figures are modified versions of image obtained from WormBase JBrowse.

A few operons were manually updated. For example, CBROP0002 and CBROPX0002 were split based on homology information in *C. elegans*, resulting in four different operons: CBROP0002A *(CBG02635, CBG02634)*, CBROP0002B *(CBG02633, CBG02632)*, CBROP0132 *(CBG01778, CBG31146, CBG01779)*, and CBROP0133 *(CBG01783, CBG01784)*. In a different case, CBROPX0007 is predicted to consist of four genes (*CBG03212*, *CBG03213*, *CBG03214*, and *CBG03215*) (Supplementary Figure 5). The *C. elegans* orthologs of these genes constitute two distinct operons (CEOP2396 and CEOP2749) (Figure 7). While the ICR between *CBG03213* and *CBG03214* is larger than 2 kb, all downstream genes in CBROPX0007 are either SL2 or SL1/SL2 spliced. Further experiments are needed to validate the structure of CBROPX0007. Table 7 lists the updated numbers of operons in each category.

**Figure 7:**
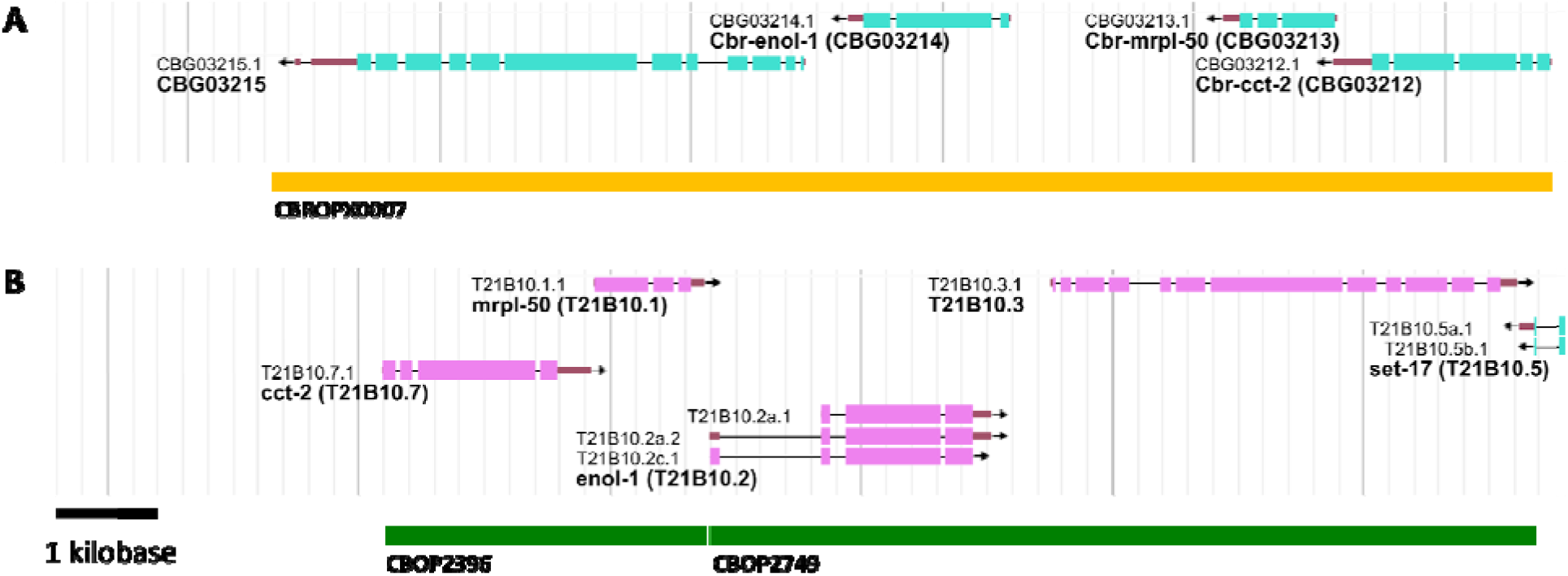
*C. briggsae* operon CBROPX0007. **(A)** A cluster of four genes that defin CBROPX0007. **(B)** The orthologs of the four genes are split between two *C. elegans* operons CEOP2396 and CEOP2749.

**Table 7:**
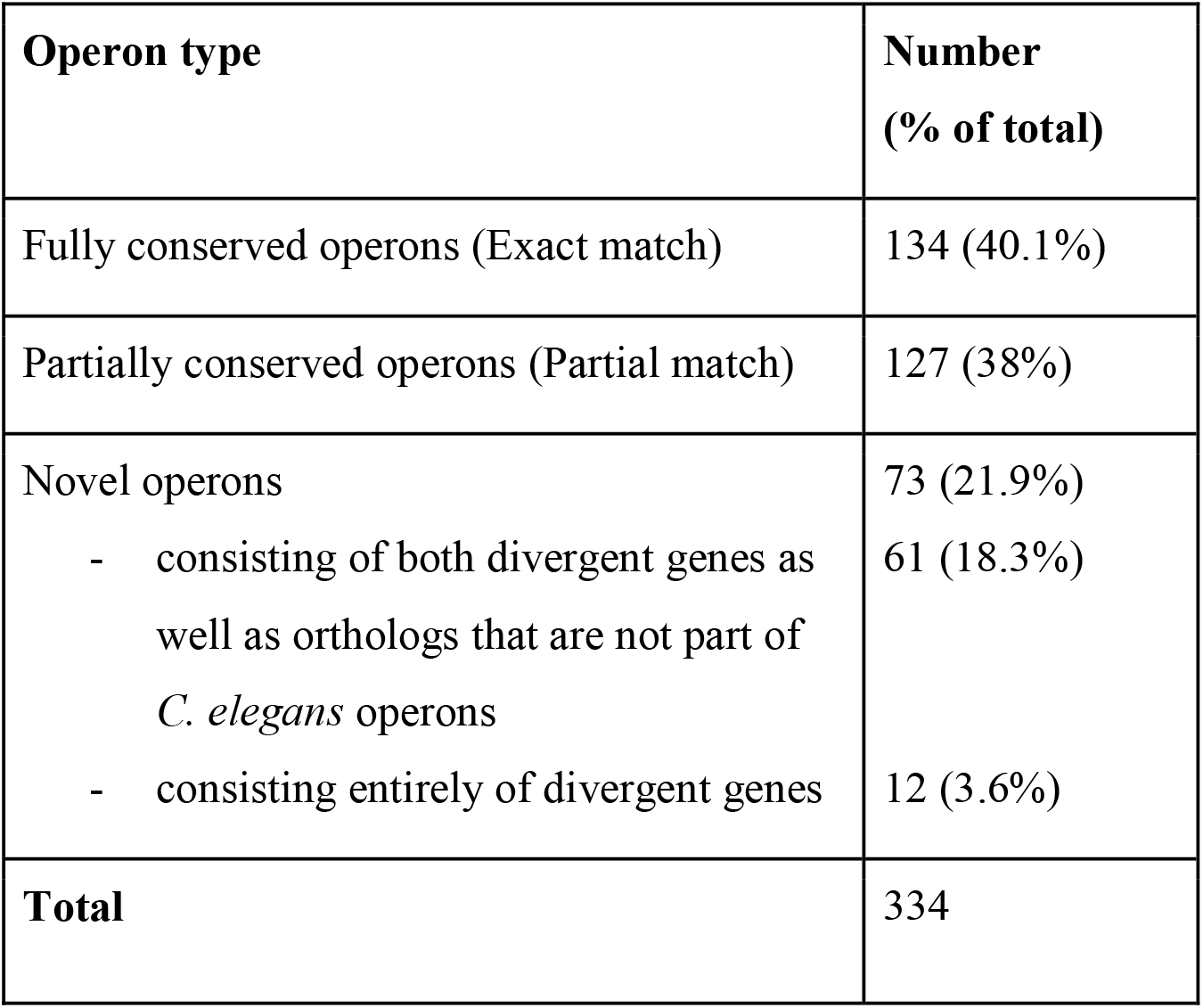
**Tag-supported operons in *C. briggsae.*** Exact match operons are conserved between *C. briggsae* and *C. elegans*. Partially conserved operons may contain some but not all orthologs that are part of corresponding *C. elegans* operons. Novel operons may contain *C. elegans* orthologs and divergent, *C. briggsae*-specific, genes.

| Operon type | Number<br>(% of total) |
| --- | --- |
| Fully conserved operons (Exact match) | 134 (40.1%) |
| Partially conserved operons (Partial match) | 127 (38%) |
| Novel operons | 73 (21.9%) |
| - consisting of both divergent genes as well as orthologs that are not part of *C. elegans* operons | 61 (18.3%) |
| - consisting entirely of divergent genes | 12 (3.6%) |
| **Total** | 334 |

We also analyzed partially conserved operons in some detail. While all of these contain *C. elegans* orthologs, their structures are not conserved. Specifically, the number of genes or some of the orthologs in corresponding operons differ between the two species (Supplementary data file 7). Of the 127 such operons, 82 contain two or more conserved genes including 58 (70% of 83) with less than 1 kb ICR between every gene. One such operon (CBROPX0003) consists of five genes (Figure 8). While the *C. elegans* operon CEOP1484 contains orthologs of all of these, CEOP1484 encompasses three additional genes.

**Figure 8.**
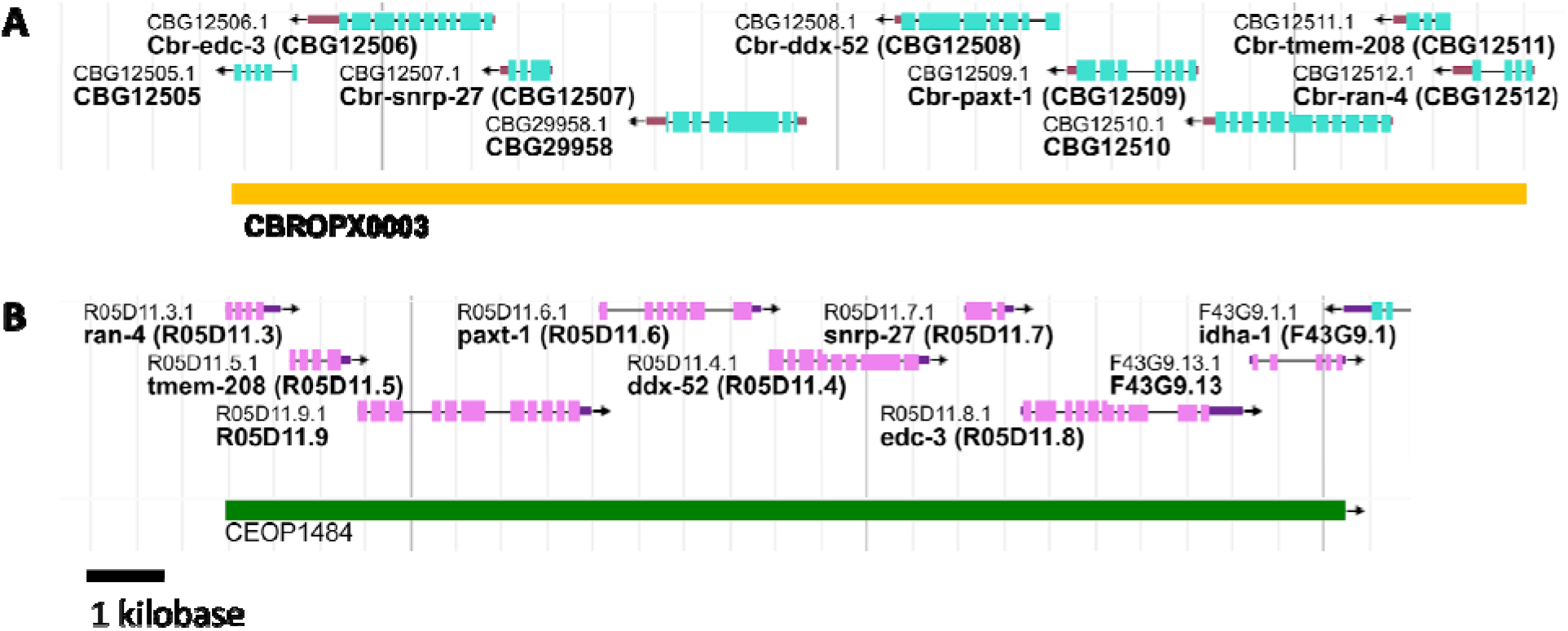
Partially conserved operon and its *C. elegans* ortholog. **(A)** CBROPX0003 is an example of a partially conserved operon identified in this study. **(B)** CEOP1484, *C. elegans* operon orthologous to *C. briggsae* operon CBROPX0003.

Our tag searches identified 73 novel operons (Supplementary data file 7). A majority of these (61, 84%) consist of a mix of conserved genes and those that lack orthologs in *C. elegans*. It is important to point out that none of the conserved genes are part of *C. elegans* operons. The other 12 (17%) operons consist entirely of genes that lack orthology in *C. elegans*. In seven of these cases, ICRs are less than 1 kb, providing further support to the operon structures (Table 8).

**Table 8:** Novel *C. briggsae* operons identified in this study with ICRs of less than 1 kb. None of the genes in these operons have orthologs in *C. elegans*. The numbers in brackets refer to ICR.

| *C. briggsae*<br>operon | No. of<br>genes | Gene names (ICR) |
| --- | --- | --- |
| CBROPX0130 | 3 | *CBG30062* (172) *CBG25686* (105) *CBG25687* |
| CBROPX0131 | 3 | *CBG27303* (533) *CBG27302* (116) *CBG27301* |
| CBROPX0140 | 2 | *CBG11551* (162) *CBG31489* |
| CBROPX0129 | 3 | *CBG21606* (235) *CBG30457* (493) *CBG21605* |
| CBROPX0139 | 2 | *CBG30329* (76) *CBG30328* |

To investigate whether the 12 novel operons are unique to *C. briggsae* or might be conserved, the analysis was extended to *C. nigoni*, a sister species to *C. briggsae* (Woodruff *et al*. 2010). For this, we manually searched 5’ upstream regions of orthologs with ≥90% sequence similarity to 5’ tags and splice acceptor sites of *C. briggsae* genes. The sequence searches revealed that four of the operons have orthologs in the same genomic order with highly similar splice site sequences and small ICRs (≥1200 bp), suggesting that they are conserved (Supplementary data file 11).

### Predicted (Non-tag-supported) operons

We report 865 predicted operons (Supplementary data file 7). While the downstream genes in these cases are spliced either with SL2 or SL1/SL2, the splicing status of the upstream gene is unknown. Most, if not all, of these are predicted to be genuine operons, especially those that are larger, i.e., consist of more than two genes. A comparison with *C. elegans* of 159 operons containing three or more genes revealed that 26 (16%) are fully conserved. A couple of examples include CBROPX0206 (5 genes) (Figure 9A, C) and CBROPX0207 (5 genes) (Figure 9D). The corresponding *C. elegans* operons are CEOP4500 (six genes) (Figure 9B, C) and CEOP5248 (seven genes) (Figure 9E). Comparison of genes in CBROPX0206 and CEOP4500 revealed that these share four orthologs. We observed two additional differences between CBROPX0206 and CEOP4500: the order of genes has changed and CBROPX0206 includes *CBG26297* which appears to lack a *C. elegans* ortholog (Figure 9C). Given that *CBG06240* and *CBG36241* are immediately upstream of CBROPX0206 and their orthologs are part of CEOP4500, the *C. briggsae* operon may be extended to include both these genes. However, we have excluded these from our operon model in the absence of corresponding TEC-RED tags. The second example, CBROPX0207, contains five genes, all of which have orthologs in CEOP5248. However, the *C. elegans* operon contains two additional genes (*ZK856.16* and *ZK856.19*) which are not conserved in *C. briggsae*.

**Figure 9:**
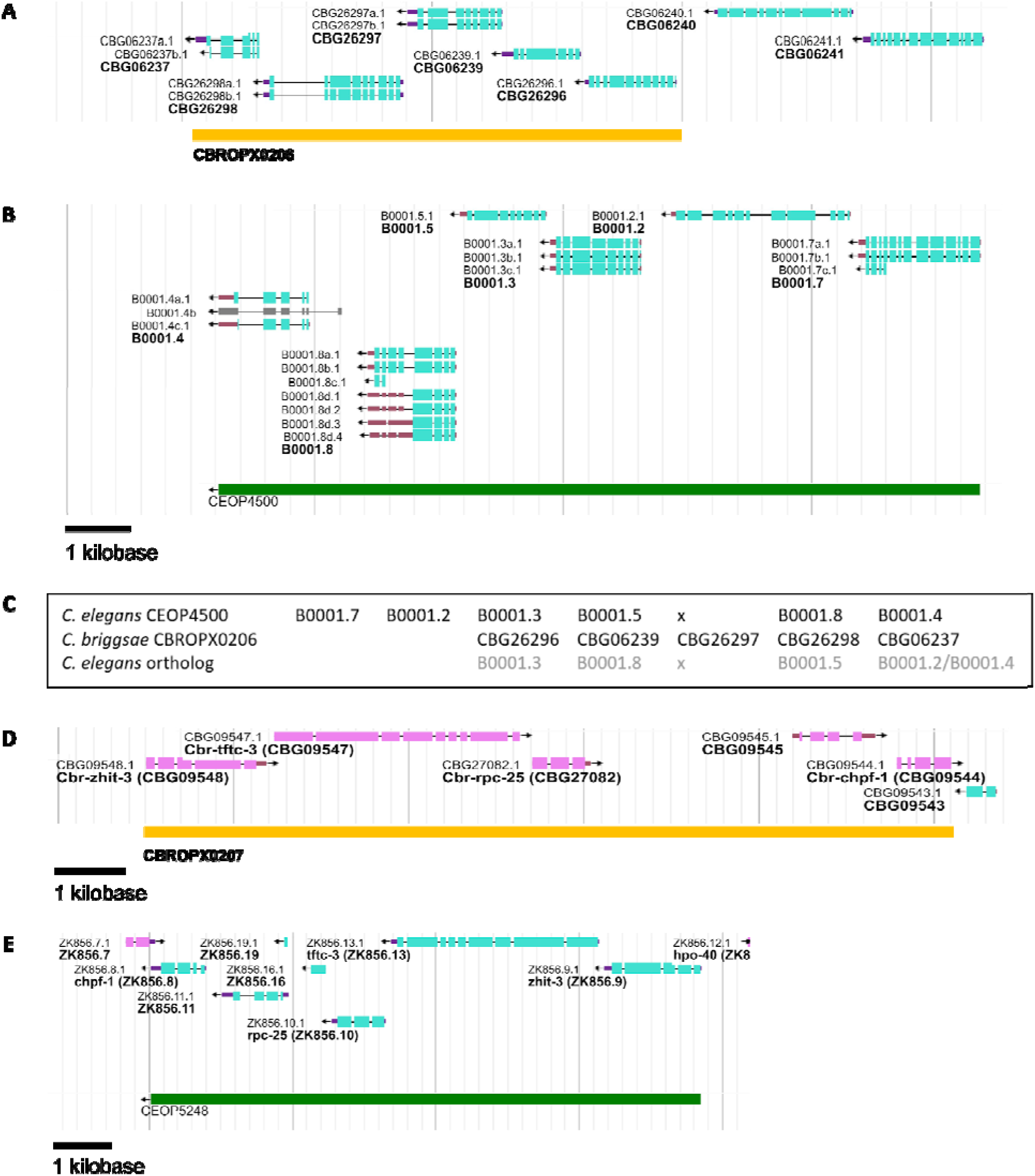
Two predicted operons in *C. briggsae* along with their *C. elegans* counterparts. **(A, B)** CBROPX0206 with five genes and its orthologous operon CEOP4500 in *C. elegans*. Three genes are conserved between these two operons. **(C)** The arrangement of genes in *C. elegans* operon CEOP4500 (row 1) and *C. briggsae* operon CBROPX0206 (row 2). The *C. elegans* orthologs of CBROPX0206 genes are shown in row 3. ‘x’ denotes a gene that is missing. *CBG06237* is orthologous to both *B0001.2* and *B0001.4*. **(D, E)** CBROPX0207 with five genes and its orthologous operon CEOP5428 with seven genes. All five genes of the *C. briggsae* operon are conserved in CEOP5428. Two additional genes are present in CEOP5428.

### SL1-type operons

We also found two operons that contain only SL1-spliced genes. These genes are positioned directly adjacent to one another, with no space between them. The SL1-type operons have been described previously in *C. elegans* and shown to lack ICR (Williams *et al*. 1999). One of the *C. briggsae* SL1-type operons consists of two genes: CBROP0134 (*CBG16825*, *Cbr-vha-11*/*CBG16826*). Its *C. elegans* ortholog, CEOP4638, also consists of two genes. Another SL1-type operon identified by our study is CBROPX0001. Its *C. elegans* ortholog is CEOP2496. Interestingly, CBROPX0001 and CEOP2496 consist of more than two genes (Figure 6). In the case of CEOP2496, the first two genes (*rpb-6* and *dohh-1*) are known to be spliced exclusively with SL1 (defined as SL1 operon) whereas the remaining downstream genes with SL2 or both SL1 and SL2.

There is also a potential SL1-type operon consisting of *CBG03984* and *CBG03983*. These two genes have a single base pair IGR (Figure 10). Interestingly, the *C. elegans* orthologs, *F23C8.6* and *F23C8.5* (SL1 and SL1/SL2 spliced, respectively) are part of one operon, CEOP1044, with an ICR of more than 400 bp (Allen et al., 2011). More work is needed to determine whether the *C. briggsae* genes are indeed part of an SL1-type operon. *C. briggsae* operons show enrichment of germline genes and highly expressed growth genes

**Figure 10.**
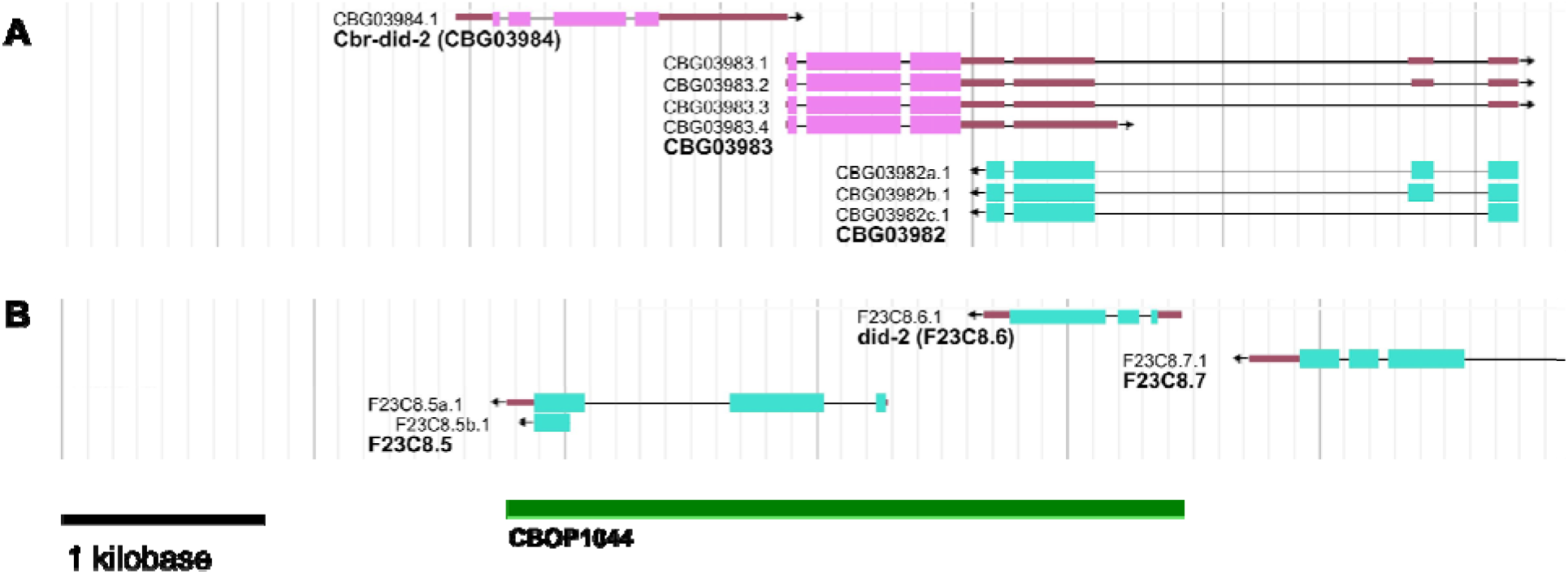
*C. briggsae* genes with a single base pair ICR. **(A)** Both *CBG03984* and *CBG03983* are spliced with SL1 leader sequences and located 1 bp apart. **(B)** *C. elegans* orthologs *did-2* and *F23C8.5*, respectively, are part of the operon CEOP1044.

Studies in *C. elegans* and *P. pacificus* have reported that germline genes are overrepresented in operons (Reinke and Cutter 2009; Sinha *et al*. 2014). We did a gene-association study in *C. briggsae* to examine a similar possibility. The results revealed a significant enrichment of germline genes in high confidence operons (p < 7.40E-98) (Supplementary data file 9).

In addition to investigating germline genes, we performed GO term analysis of operon genes and found enrichment of terms associated with metabolic and biosynthesis processes. The pattern of enrichment was similar to what was observed with a *C. elegans* operon dataset (Supplementary data file 9). We also found enrichment of growth-related genes, as in *C. elegans*, specifically, female gamete generation (GO:0007292), embryo development ending in birth or egg hatching (GO:0009792), reproduction (GO:0000003) and embryo development (GO:0009790) (Zaslaver *et al*. 2011). It is important to point out that while GO terms are similar in both species, *C. briggsae* operon genes associated with specific processes are not always the orthologs of *C. elegans* gene sets. We therefore conclude that functions of operon genes are conserved even if specific genes are not.

## DISCUSSION

This paper reports major improvements in the genome annotation of *C. briggsae*. We recovered 10,243 unique 5’ end tags with matches in the genome, of which 6,395 correspond to SL1 and SL2 spliced exons and provide support to the existence of 4,252 unique *trans*-spliced genes. Another 362 genes have been identified as paralogs, including 42 for which the paralogous relationship is supported by WormBase annotation. We also report 52 novel exons that may define new genes or exons of existing genes. Figure 11 provides a global overview of sequences identified in our analysis.

**Figure 11.**
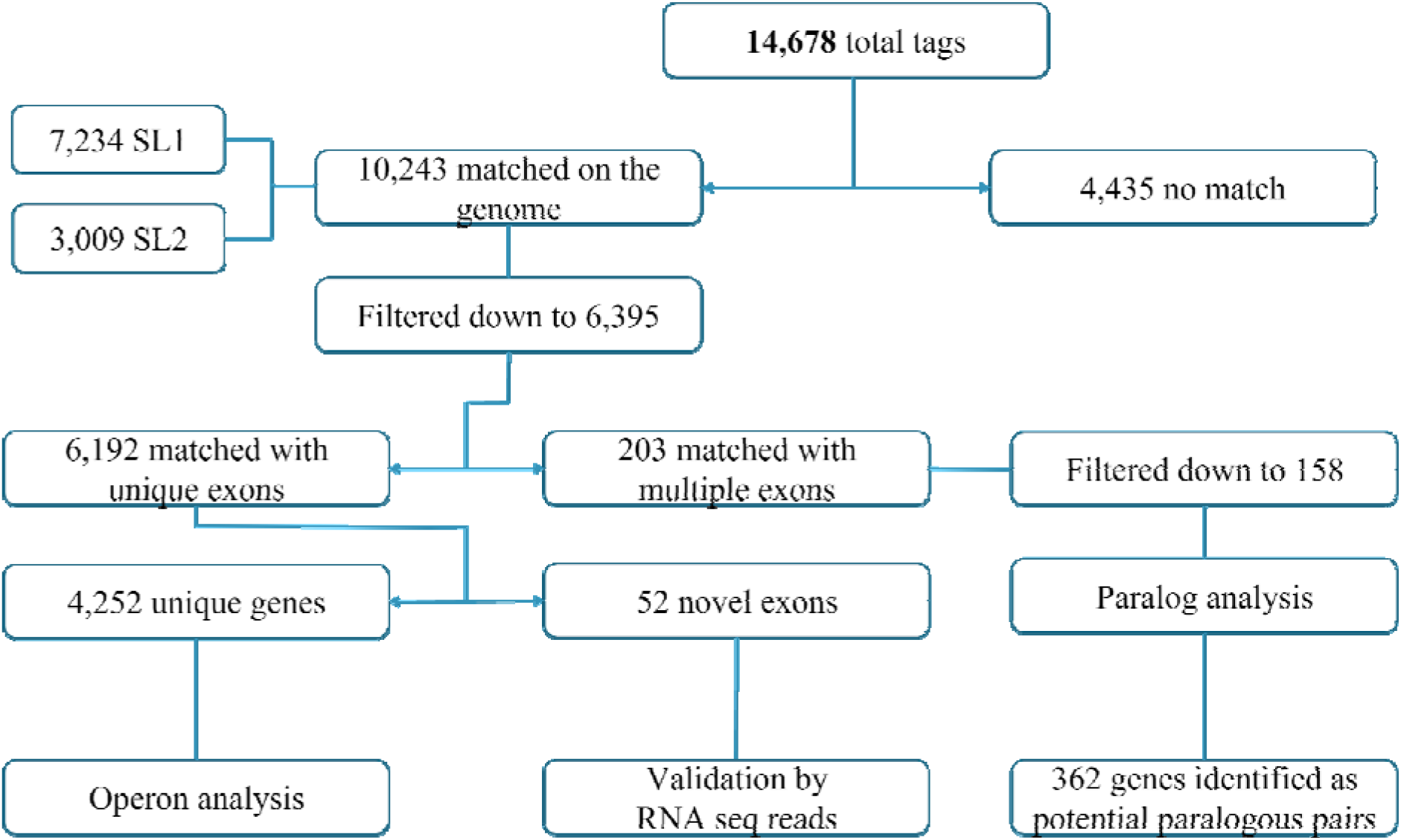
An overview of TEC-RED analysis in *C. briggsae*. The 5’ sequence tags were used to identify exons and genes. Further analysis resulted in the discovery of operons, paralogs, and novel exons.

In *C. elegans,* 84% of all genes are spliced to leader sequences (Tourasse *et al*. 2017). If the percentage is comparable in *C. briggsae*, then our work has resulted in the identification of roughly one-quarter of all *trans*-spliced genes in this species. Further analysis has revealed that two-thirds of all *C. briggsae* genes are spliced with SL1, while the rest are split evenly between SL2 and SL1/SL2 hybrid sequences (65% SL1, 18% SL2 and 18% SL1/SL2). Assuming that the TEC-RED method is unbiased in regard to the recovery of SL1 and SL2 spliced transcript tags, the proportion of spliced genes in *C. briggsae* differs from those in *C. elegans* as reported by Allen et al. (Allen *et al*. 2011), (82% SL1, 12% SL2 and 8% SL1/SL2). Additionally, 14 genes were found to be spliced to leader sequences only in *C. briggsae* and not in *C. elegans*. More work is needed to determine if *trans*-splicing of these genes has indeed diverged between the two species.

Our analysis revealed that most of the genes identified by unique tag matches are represented by a single transcript (94.8%) and very few (5.2%) by multiple transcripts. Studies in *C. elegans* have reported roughly 18% of genes giving rise to multiple isoforms (Wang *et al*. 2010; Spieth *et al*. 2014), although this number is predicted to be as high as 25% (Ramani *et al*. 2011; Zahler 2012). Considering this, along with the fact that our experiments captured only a partial set of all spliced genes, the actual proportion of genes with multiple transcripts in *C. briggsae* is likely to be much higher. Among other things, it was found that 77.8% of the genes in our study have 5’ start sites that match with those annotated by WormBase. The remaining ones were considered mispredictions, most of which were major mispredictions (15.7%) as 5’ start sites in these cases map anywhere between 20 bp to 3 kb away from known locations. We also found 52 new, previously unreported exons that map more than 3 kb upstream to the nearest exon of existing genes, and potentially include some that define 5’ start sites of new genes.

Several approaches were taken to validate tag-based gene models. One involved comparing results with those obtained using an older genome annotation (gff) file. The findings revealed that a total of 858 genes for which 5’ ends were correctly annotated in WS276 were mispredicted or absent in the older version, which demonstrates that our data can help improve start sites of many *C. briggsae* genes. Another approach involved comparing 5’ ends of some of the genes with those of *C. elegans* orthologs. Of the 21 alternate start sites and 50 major mispredicted start sites analyzed, 38% and 34%, respectively, are supported by *C. elegans* transcripts. Finally, we examined the 52 newly discovered exons and found that 37% of these are supported by RNA-seq data in WormBase. The above three validations provide significant support to the accuracy of our analysis of expressed transcripts in *C. briggsae*.

The identification of genes spliced with leader sequences in *C. briggsae* allowed us to curate operons and study their conservation. Even though the operon-based organization of genes in *C. elegans* and *C. briggsae* is similar to those found in bacteria and archaea, work in *C. elegans* has shown that worm operons have no ancestral relationship with prokaryotes and appear to have evolved independently within the nematode phylum (Blumenthal 2004; Qian and Zhang 2008). We identified a total of 1,199 operons, of which 28% consist entirely of tag-supported genes. Of the remaining operons with partial tag support, 159 contain three or more genes.

Combined with the fully tag-supported operons, this totals to 493 operons in *C. briggsae* with a high degree of confidence. Comparison of tag-supported operons with *C. elegans* revealed that 134 (40%) are conserved, with the remainder being partially conserved (127, 39%) and novel (73, 21%). A subset of novel operons (12, 17%) consists entirely of genes that lack *C. elegans* orthologs. Further comparisons with *C. nigoni* revealed that four of the twelve are likely to be conserved, suggesting that these might have arisen in the common ancestor shared between *C. briggsae* and *C. nigoni*. Whether the remaining eight are unique to *C. briggsae* requires more analysis. Along with the above-mentioned operons, we also uncovered two conserved SL1-type operons. Together, these data demonstrate that while many of the operons are conserved, there are substantial differences between the two species. The findings represent the first comprehensive analysis of operons in *C. briggsae*.

In conclusion, the data presented in this study has significantly improved the annotation of the *C. briggsae* genome by validating existing gene models, refining start sites of many genes, identifying novel gene exons, alternate transcripts, and by providing a comprehensive analysis of operons and paralogous gene sets. While the majority of the genes and operons are conserved in *C. elegans*, our work has also revealed substantial differences between the two species. The improvements to the genome annotation reported here are expected to strengthen *C. briggsae* as a model for comparative and evolutionary studies.

## DATA AVAILABILITY

The data underlying this article are available in the article and in its online supplementary material.

## ACKNOWLEDGEMENTS

We thank WormBase for assistance with some aspects of data analysis, Mary Ann Allen and Tom Blumenthal for discussions on *C. elegans* operons, and members of the Gupta lab for feedback on the manuscript. This work was supported by grants to BPG (NSERC Discovery) and PWS (U24-HG002223). PWS was an Investigator with the HHMI, which partially supported this work.

## SUPPLEMENTARY MATERIALS

## SUPPLEMENTARY DATA FILES (Microsoft Excel spreadsheets)

| File name | Description |
| --- | --- |
| Supplementary data file 1 | Exons identified in our analysis |
| Supplementary data file 2 | Unique genes identified |
| Supplementary data file 3 | Genes uniquely spliced in *C. briggsae* |
| Supplementary data file 4 | Validation based on overlap with WS176 |
| Supplementary data file 5 | New exons identified by our study |
| Supplementary data file 6 | Manual curation of 1b and major mispredictions based on *C. elegans* orthologs |
| Supplementary data file 7 | List of operons |
| Supplementary data file 8 | Intergenic region values |
| Supplementary data file 9 | Germline genes present in operons and GO analysis |
| Supplementary data file 10 | Proposed paralog sets |
| Supplementary data file 11 | Matches to *C. briggsae* novel operons in *C. nigoni* |

## SUPPLEMENTARY TABLES

**Supplementary Table 1:**
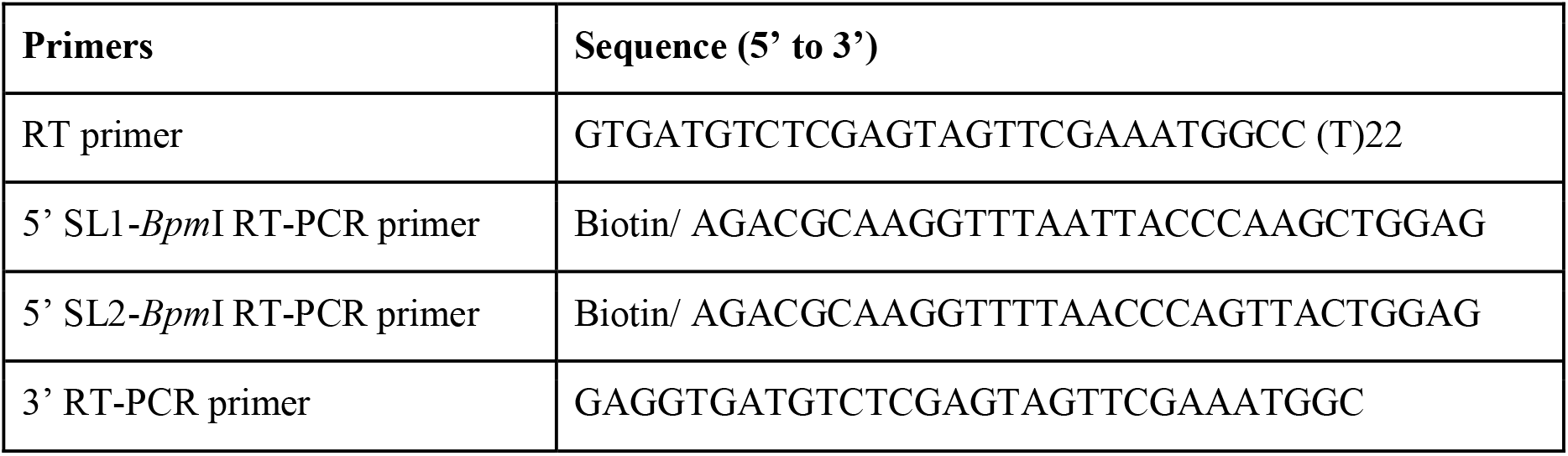
Primers used to generate Biotin-RT-PCR products

**Supplementary Table 2:**
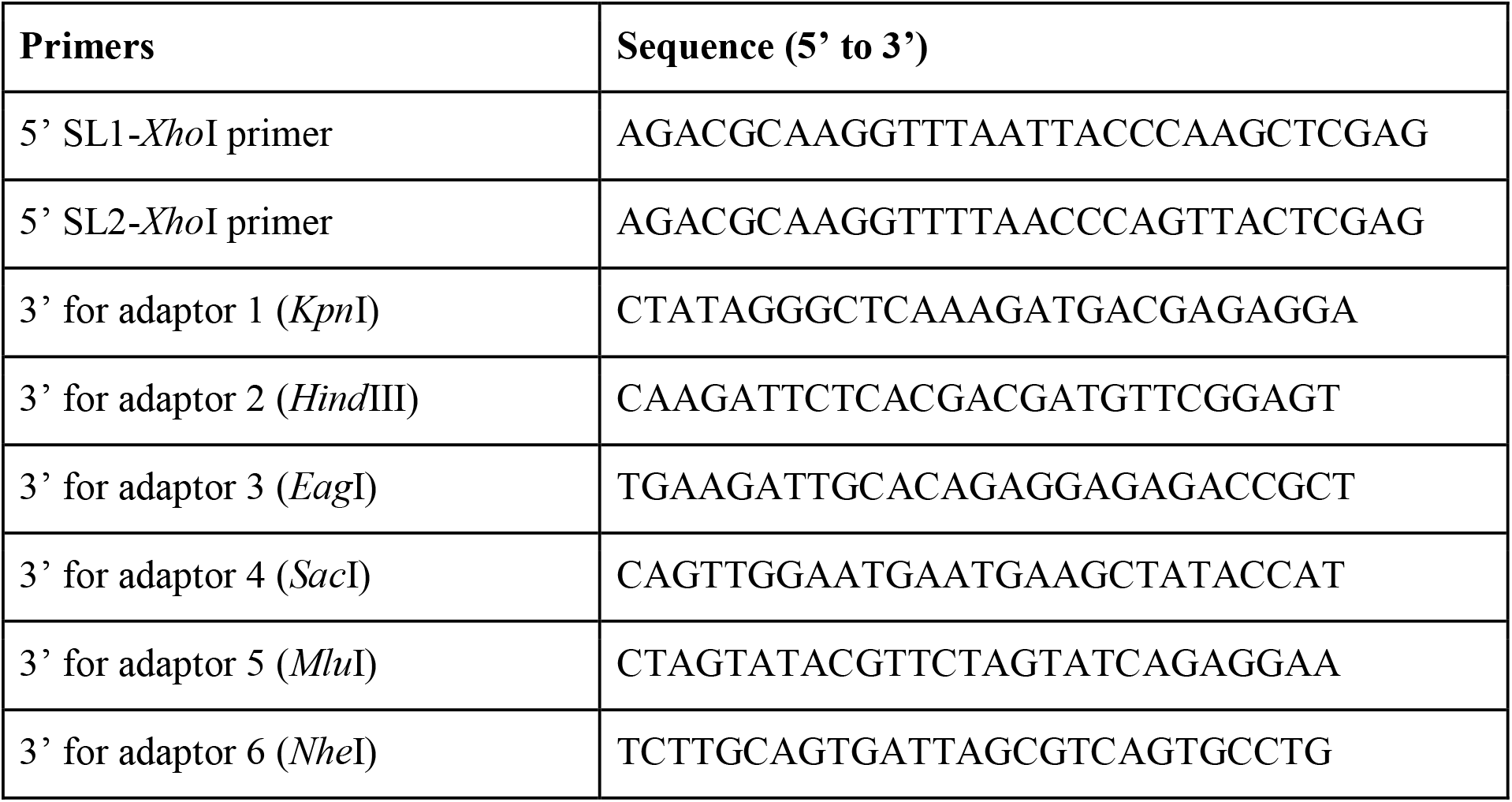
PCR primers used to generate mono-tags from the 5’ biotin-adaptor DNA fragments

**Supplementary Table 3:**
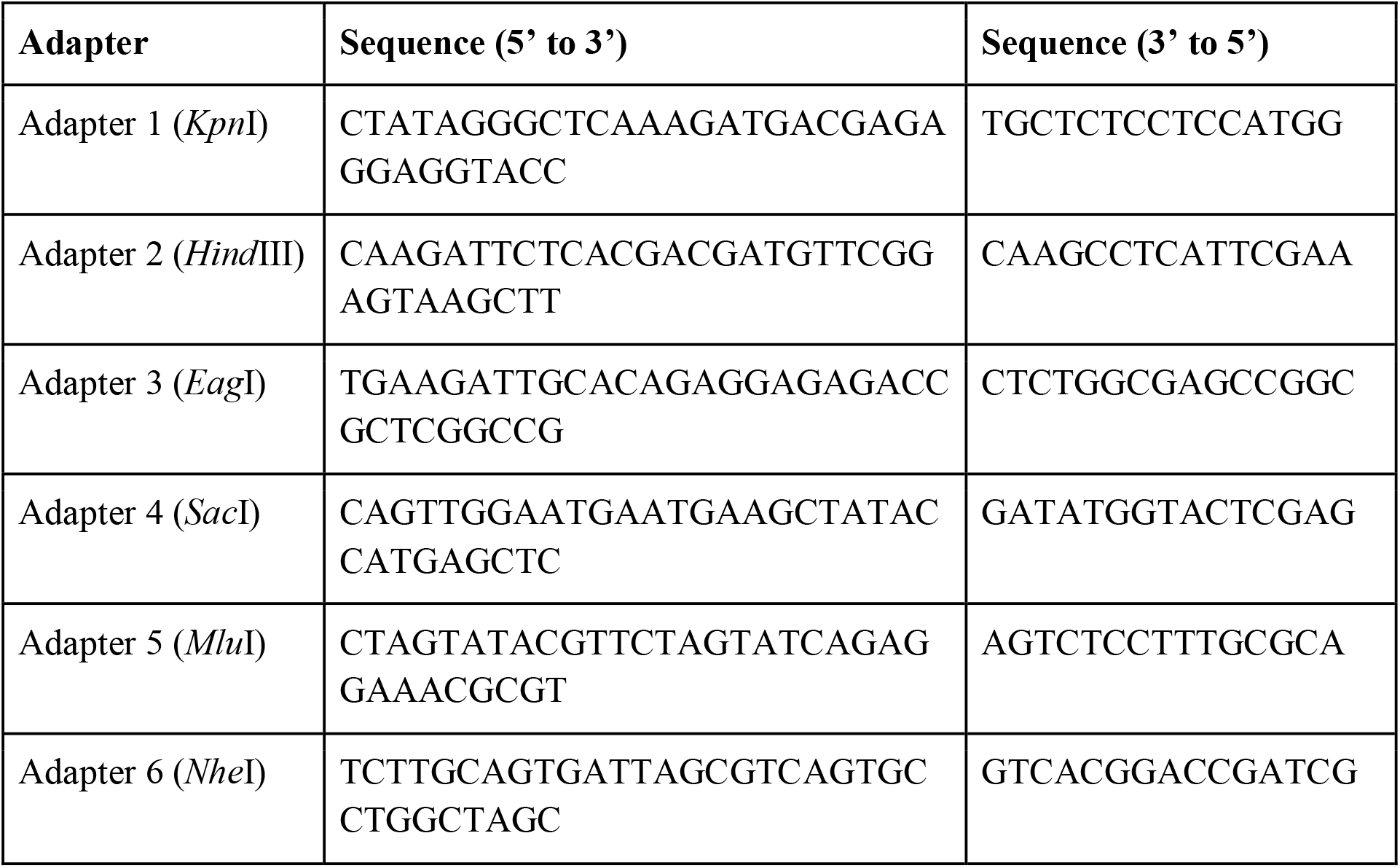
Adaptors used for ligation onto *Bpm*I-digested, 5’ biotin-DNA fragments

**Supplementary Table 4:** Chromosomal locations of 4,252 unique genes identified by TEC-RED. Chr: Chromosome, Un: unmapped genomic region.

| Chr | Total<br>gene<br>count | SL1<br>genes | Fraction | Density | SL2<br>genes | Fraction | Density | SL1/SL2<br>genes | Fraction | Density | Chr length<br>(Mb)* |
| --- | --- | --- | --- | --- | --- | --- | --- | --- | --- | --- | --- |
| I | 763 | 447 | 16.26 | 28.93 | 158 | 21.27 | 10.23 | 158 | 20.79 | 10.23 | 15.45 |
| II | 752 | 473 | 17.21 | 28.46 | 147 | 19.78 | 8.84 | 132 | 17.37 | 7.94 | 16.62 |
| III | 751 | 444 | 16.15 | 30.47 | 145 | 19.52 | 9.95 | 162 | 21.32 | 11.12 | 14.57 |
| IV | 717 | 447 | 16.26 | 25.57 | 133 | 17.90 | 7.61 | 137 | 18.03 | 7.84 | 17.48 |
| V | 749 | 534 | 19.43 | 27.40 | 105 | 14.13 | 5.39 | 110 | 14.47 | 5.64 | 19.49 |
| X | 515 | 400 | 14.55 | 18.57 | 54 | 7.27 | 2.51 | 61 | 8.03 | 2.83 | 21.54 |
| Un | 5 | 4 |  |  | 1 |  |  | 0 |  |  |  |
|  | 4252 | 2749 |  | 26.14 | 743 |  | 7.07 | 760 |  | 7.23 | 105.15 |

\*Ross, J. A., D. C. Koboldt, J. E. Staisch, H. M. Chamberlin, B. P. Gupta et al., 2011 *Caenorhabditis briggsae* recombinant inbred line genotypes reveal inter-strain incompatibility and the evolution of recombination. PLoS Genet 7: e1002174.

**Supplementary Table 5:** Intergenic distances of selected Category 3 genes that are less than 10 kb apart.

| Adjacent genes identified by tags | IGR | BLAST alignment | *C. elegans* orthologs | *C. briggsae* gene orientation |
| --- | --- | --- | --- | --- |
| *CBG25816*,<br>*CBG00473* | 2,903 bp | Yes | None | Opposite |
| *CBG08766*,<br>*CBG08768a* | 578 bp | No<sup>*</sup> | *F25E5.8* and *nhr-117* | Opposite |
| *CBG25203*,<br>*CBG29819* | 2,251 bp | No | *F59A3.2* and *ubl-5* | Opposite |
| *CBG26374*,<br>*CBG05421* | 3,564 bp | No<sup>*</sup> | None, *fan-1* | Same |
| *CBG26845*,<br>*CBG26846* | 8,559 bp | Yes | None | Same |

<sup>\*</sup>BLAST match showed some similarity in a very small 5' region.

## SUPPLEMENTARY FIGURES

**Supplementary Figure 1:**
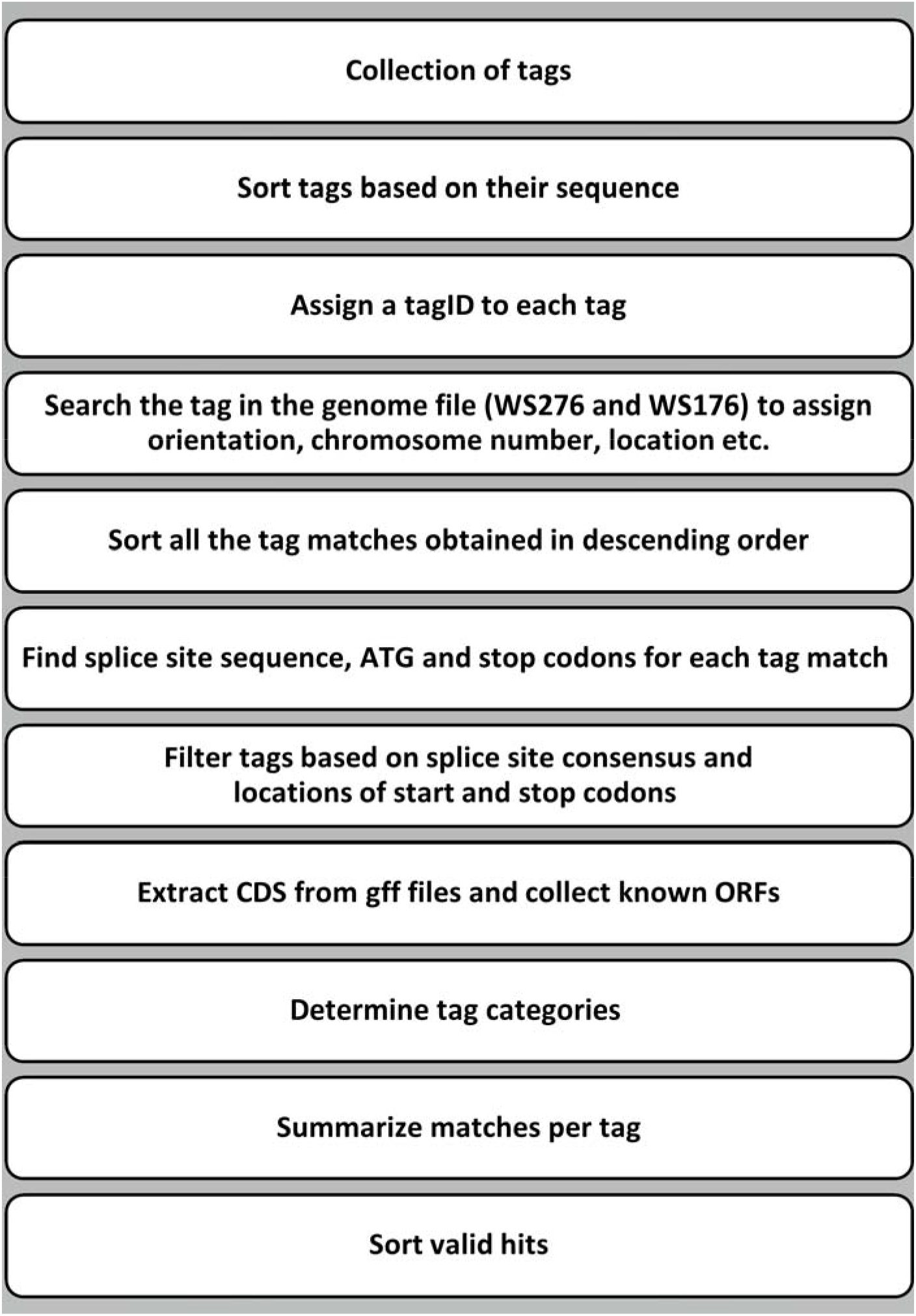
Flowchart of steps used to analyze 5’ tag sequences and genes.

**Supplementary Figure 2:**
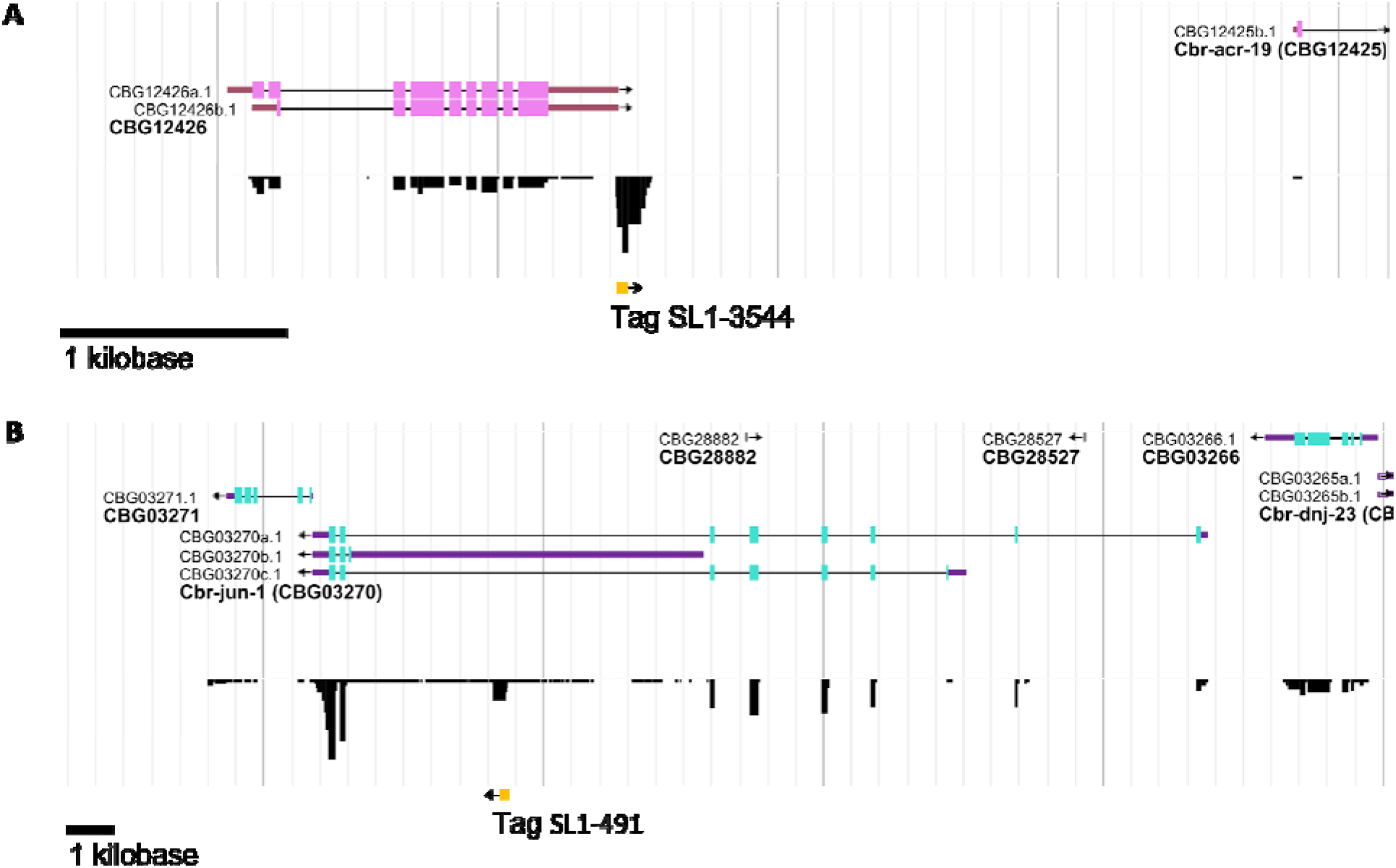
Selected examples of novel exons identified by tags that are supported by WormBase RNASeq data. The top track of the genome browser shows currently curated genes. Second track shows alignments of short read sequences from all available RNASeq projects on WormBase. The number of reads has been normalized by averaging over the number of libraries. The height of reads boxes indicates the relative score of the feature. The bottom track shows a TEC-RED tag binding at a genome location predicted to contain the 5’ start site of a new exon. **(A)** New exon between *CBG12426b.1* and *CBG12425b.1*. **(B)** A new exon inside *CBG03270a.1*.

**Supplementary Figure 3:**
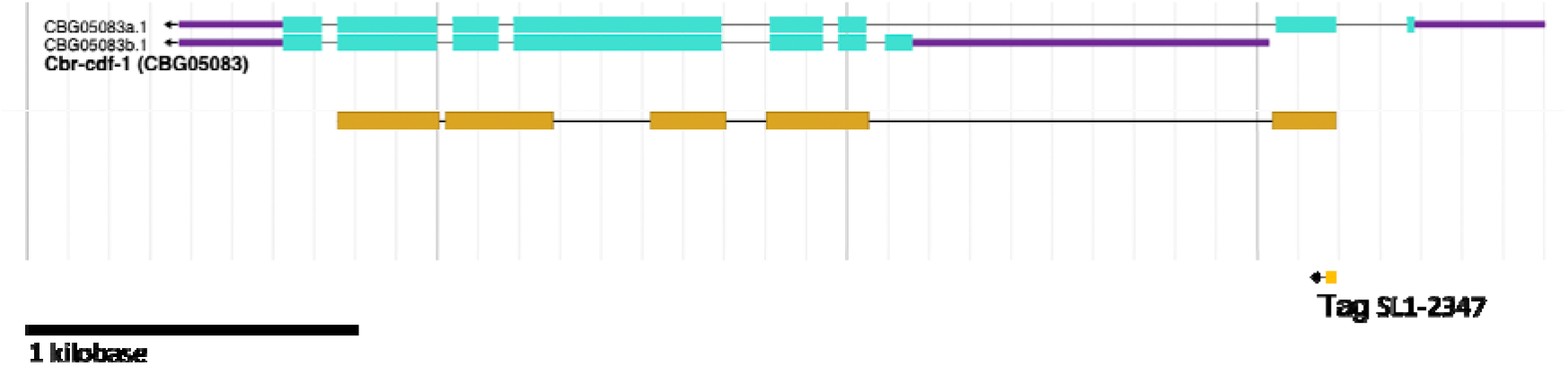
An example of the 1b category in *Cbr-cdf-1* in WormBase genome browser. The top track shows curated gene *Cbr-cdf-1*. The middle track shows the *C. elegan C15B12.7a.1* (*cdf-1*) gene model, which is indicated in orange. The bottom track shows a category 1b TEC-RED tag binding at the 5’ start site of exon 2 of *Cbr-cdf-1*. The *C. elegan* gene model supports the 5’ start site of a new transcript variant for *Cbr-cdf-1*.

**Supplementary Figure 4.**
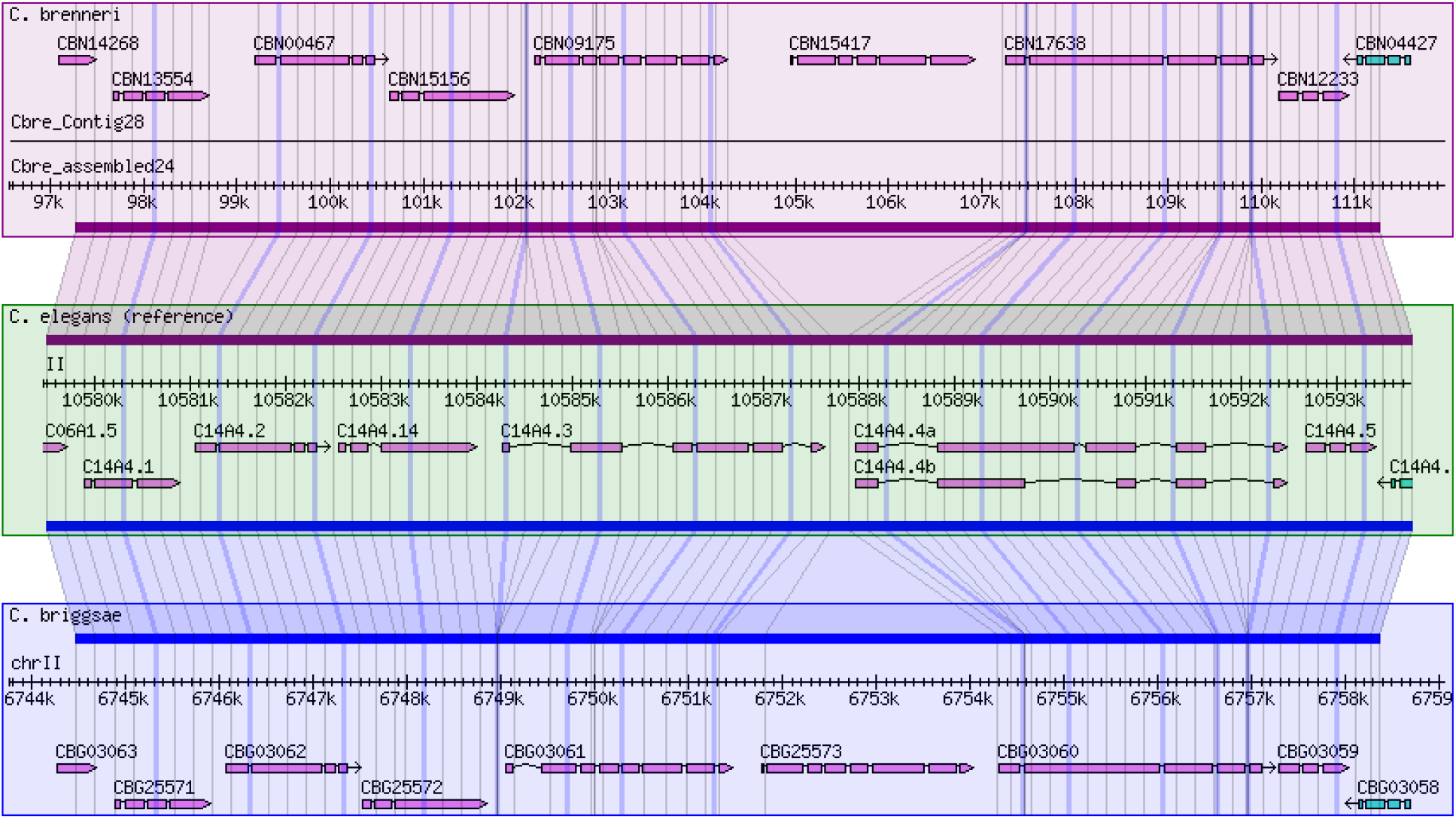
A screenshot of the WormBase synteny browser showing th genomic region of *C. briggsae* operon CBROPX0001, and the corresponding regions in *C. brenneri*, *C. elegans* and *C. briggsae*.

**Supplementary figure 5:**
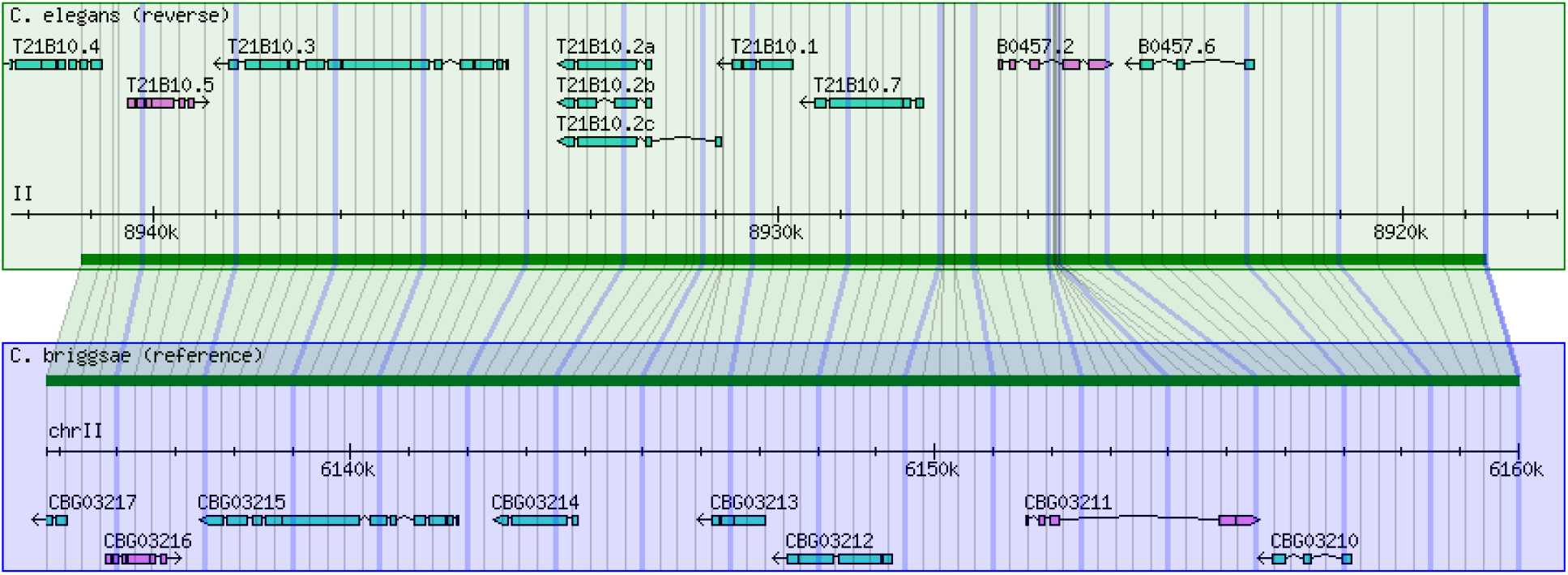
A screenshot of the WormBase synteny browser showing the genomic region consisting of a cluster of four *C. briggsae* genes that define the CBROPX0007 operon and corresponding region in *C. elegans*.

